# Effects of Satellite-Linked Telemetry Tags on Humpback Whales in the Gulf of Maine: Photographic Assessment of Tag Sites

**DOI:** 10.1101/2024.02.07.579298

**Authors:** Frances M.D. Gulland, Jooke Robbins, Alexandre N. Zerbini, Virginia Andrews-Goff, Martine Bérubé, Phillip J. Clapham, Michael Double, Nicholas Gales, Amy S. Kennedy, Scott Landry, David K. Mattila, Doug Sandilands, Jennifer E. Tackaberry, Marcela Uhart, Ralph E. T. Vanstreels

**Affiliations:** University of California, Davis, CA, USA; Center for Coastal Studies, Provincetown, MA, USA; Cooperative Institute for Climate, Ocean and Ecosystem Studies, University of Washington, Seattle, WA, USA; Marine Mammal Laboratory, Alaska Fisheries Science Center, NOAA, Seattle, WA, USA; Marine Ecology and Telemetry Research, Seabeck, WA, USA; Cascadia Research Collective, Olympia, WA, USA; Australian Antarctic Division, Hobart, Tasmania, Australia; Marine Evolution and Conservation, Groningen Institute of Evolutionary Life Sciences, University of Groningen, Groningen, The Netherlands; Seastar Scientific, Vashon Island, WA, USA

**Keywords:** HEALTH, MODELLING, PHYSIOLOGY, SATELLITE TAGGING, TELEMETRY

## Abstract

Hundreds of large whales have been tracked using consolidated (Type-C) satellite tags, yet there have been few studies on their impacts on whale health. In 2011, we initiated the first study designed to evaluate the effects of these tags in a baleen whale. Between 2011 and 2018, we tagged 79 North Atlantic humpback whales in the Gulf of Maine. We initially deployed commonly-used tags with an articulation between the anchor and transmitter (n=35, 2011-2012). However, evidence of breakage prompted the development and use of more robust, integrated tags (n=45). Tagged individuals were photographed immediately prior to, during and up to 11 years after tagging. They were re-encountered on an average of 41.3 days (SD=44.3), yielding 2,971 photographed sightings through 2022. An objective scoring system was developed to characterise tag site tissue responses based on photographs and to identify risk factors for prolonged healing. The initial tissue response to tagging was minimal, followed by skin loss around the tag, sometimes a degree of swelling, occasional extrusion of blubber, changes in skin colour, local depression formation, tag loss and skin healing over the tag site, sometimes with a depression remaining. At last sighting, most non-integrated and integrated tag sites exhibited small shallow skin depressions (58.8% and 66.7%, respectively). Some exhibited deeper depressions with differing adjacent skin coloration (26.5% and 15.6%, respectively) or barely detectable marks (11.8% and 15.6%, respectively). Mild swellings occasionally persisted at the tag site, but this was uncommon for both tag designs (2.9% and 2.2%, respectively). More severe tissue responses were associated with non-integrated tags and placements lower on the body. This study highlights the importance of using robust tag designs to minimise negative effects from Type-C tags. Furthermore, because tag placement was shown to affect outcome, precision equipment, experienced taggers and vessel operators are critical for optimal deployments.

## INTRODUCTION

Monitoring the location and movements of whales can be vital to research and conservation (Hays *et al*., 2019). One approach for collecting such data is the attachment of instruments that record and transmit information, commonly referred to as “tags”. In the case of baleen whales, the only tags that are consistently capable of collecting data over weeks to months (rather than hours to days) consolidate their electronics and retention elements into a single unit that is embedded in the body of the whale. These “Type-C” tags have been deployed on hundreds of whales, including humpback (*Megaptera novaeangliae*), bowhead (*Balaena mysticetus*), fin (*Balaenoptera physalus*), blue (*Balaenoptera musculus*), North Atlantic, North Pacific and southern right (*Eubalaena glacialis, E. japonica, and E. australis*), gray (*Eschrichtius robustus*), minke (*Balaenoptera acutorostrata*) and sperm whales (*Physeter macrocephalus*) (e.g., Baumgartner & Mate 2005; Zerbini *et al*. 2006; Mate *et al*. 2007; Gales *et al*. 2009a; Best *et al*. 2015; Zerbini *et al*. 2015; Sepúlveda *et al*. 2018; Willson *et al*. 2018; Zerbini 2018). However, Type-C tags are designed to anchor in the connective tissue underlying the blubber or into the muscle (Mate *et al*. 1983; Andrews *et al*. 2019) and the associated wounds have the potential to cause pain and discomfort. This potential for harm is important both because it could affect the data collected and because of animal welfare and conservation concerns.

Potential impacts of tagging include effects on whale behaviour, health, reproduction and survival (Weller 2008). Limited repeated sightings of tagged whales, small sample sizes, lack of data from control (un-tagged) whales from the same study populations, and annual variability in reproductive rate and survival challenge the determination of such impacts (Robbins *et al.,* in prep). There have been few studies on the impacts of tags on cetaceans, and a paucity of information on tissue responses and healing of the tag placement wound, mostly because dedicated effort to conduct follow-up studies has been minimal (Weller 2008; ONR 2009; Andrews *et al*. 2019). Prior attempts were based primarily on opportunistic observations of previously tagged fin and humpback whales in Alaska (Watkins *et al*. 1981; Mizroch *et al*. 2011), North Atlantic right whales (Kraus *et al*. 2000), southern right whales (Best & Mate 2007) and various other whale species tagged by Mate and colleagues (Mate *et al*. 2007; Gendron *et al*. 2015; Norman *et al*. 2018). Evaluation of the potential impacts of a tag on whale health can include systematic assessment of health indicators, such as the animal’s behaviour, nutritional status, skin condition, hormonal status and parasitic load, as well as assessment of the localised tissue changes at the tag attachment site. As future studies using tags are developed to inform research and guide conservation, it is essential that there be concurrent evaluation of the impacts of these studies on individuals. Such work has been recommended by the ethical guidelines for scientists from the Society for Marine Mammalogy (Gales *et al*. 2009b) and is critical for informed decisions that balance individual animal welfare with population conservation (Robbins *et al*. 2013; Willson *et al*. 2018; Andrews *et al*. 2019; Papastavrou & Ryan 2023).

For some species, standardised photographic methods have been developed for assessing nutritional and general health (Pettis *et al*. 2004; Bradford *et al*. 2012; Christiansen *et al*. 2019; Hörbst 2019), as well as injuries from entanglements (e.g., Robbins & Mattila 2004; Knowlton *et al*. 2012), ship strikes (Hill *et al*. 2017) and predation (e.g., Naessig & Lanyon 2004; Mehta *et al*. 2007; Steiger *et al*. 2008). This contrasts with the lack of standardised objective criteria for assessment of wounds associated with satellite tagging. Assessment of the impact of a foreign body or wound in a mammal classically involves use of hands-on physical examinations, imaging of tissues using radiology, ultrasound or infra-red technology, and biopsy for histology and microbiology, ideally at serial intervals (Gulland *et al*. 2018). In living, free-ranging whales, examination to date has been limited to photographic assessments of tag sites, often at a single time point, with qualitative, opinion-based, assessments of wound severity (Norman *et al*. 2018).

In 2011, we initiated the first study designed to evaluate the range of potential effects of Type-C tags on humpback whales while improving understanding of their seasonal movements and residency patterns. The project was envisioned and implemented as a collaboration among scientists with extensive experience in large whale biology, behaviour, photo-identification, tagging, health assessments and veterinary medicine. It was undertaken with a goal of transparent communication with other scientists, resource managers and permitting agencies. As part of this overarching project, a scoring system was developed for evaluating tag site wounds based on photographs (Robbins *et al*. 2013; Andrews *et al*. 2019). This system has already been used for evaluating tag site wounds on Arabian Sea humpback whales (Minton *et al*. 2022) and southern right whales (Charlton *et al*. 2023). Here we further refine this method and report on its use to characterise the sequence of tissue responses to tagging over a decade in one well-studied population of North Atlantic humpback whales in the Gulf of Maine (GoM). We provide data on the effects of a widely used tag that exhibited a high incidence of breakage, as well as of improved, robust tags that performed as designed.

## MATERIALS AND METHODS

### Gulf of Maine humpback whales

This study focused on humpback whales in the GoM, which arrive on their feeding grounds as early as mid-March and persist in the region through December (Clapham *et al*. 1993; Robbins 2007). Individual humpback whales were identified in the field based on their natural markings (Katona & Whitehead 1981), and many animals in this population have extensive sighting and demographic data accumulated over decades. Individuals were selected for tagging by a population expert (JR) based on life history and demographic traits documented in a four-decade GoM humpback whale catalogue curated by the Center for Coastal Studies (CCS, Provincetown, MA). Individuals with the following characteristics were prioritised for inclusion in the study: 1) regular annual return to the primary study area, Stellwagen Bank in the southwestern GoM, 2) frequent within-season re-sighting rates, 3) known sex and age class as well as detailed calving history (in the case of females), 4) apparent good skin health and nutritional status at time of tagging, and no recent history of poor health. Candidates were also selected with a goal of a balanced sample of males and females. The sex of individuals was known from prior genetic analysis of a biopsy sample (Palsbøll *et al*. 1992; Bérubé & Palsbøll 1996a, b)and typically further supported by prior documentation of the genital slit (Glockner 1983) and/or calving history in the case of females. Finally, the project focused on adults without a dependent calf in the tagging year to maximise sample sizes, and to minimise survival effects given that juveniles and lactating females were known to have lower annual probabilities of survival (Robbins 2007). Age class was assigned based on the number of elapsed years since the first sighting, which was based either on the year of birth (exact age) or a subsequent sighting after independence (minimum age). Individuals were considered to be juveniles if younger than the earliest known age at parturition in this population (5 years, Clapham 1992).

### Satellite tag deployments

Between 2011 and 2018, 79 humpback whales were tagged under permits issued by the U.S. National Marine Fisheries Service under the U.S. Endangered Species Act and the Marine Mammal Protection Act. This research was also reviewed for ethical considerations and to conform to the U.S. Animal Welfare Act by the Internal Animal Care and Use Committee from NOAA’s Marine Mammal Laboratory (Seattle, WA).

All of the tags were designed to penetrate skin and blubber and anchor in or below the fascia between the blubber and muscle, with tags of varying tag lengths (between 28 and 30 cm). Five different designs were used, starting with a tag described by Gales *et al*. (2009a). Each subsequent tag version reflected one or more improvements to improve the design based on information provided by this study, as described in detail by Zerbini *et al*. (in prep). The tags deployed in 2011 and 2012 were designed with an articulating anchor that was intended, in part, to reduce injury due to shearing between the blubber and muscle layers (Moore *et al*. 2013; Moore & Zerbini 2017). However, the present study revealed that structural interfaces in the anchoring system, or between the anchor and the electronics package, could not withstand the forces on deployment and/or as experienced while deployed in the whale. This resulted in tag breakage on or after deployment. The tag design was subsequently changed to increase its robustness by developing fully integrated tag designs in 2013, 2015 and 2018 (Zerbini *et al*., in prep.). We therefore differentiate between “non-integrated” tags and the improved “integrated” tag designs that had no flexible tag parts and more robust interfaces.

Regardless of design, all external tag components were built from surgical quality stainless steel. Each tag was sterilised with ethylene oxide gas and stored in a sealed autoclave pouch until the time of deployment. Tag deployments were performed from the bow of a 12.1 m twin-screw motor vessel or from a raised platform on a 10.7 m, dual outboard, rigid-hulled boat. Both vessels were equipped with top-side steering, either in the form of a tower or flying bridge. Approaches were made at distances ranging from 3-10-m from the whale by a single, expert vessel operator (DKM). Tags were deployed by experienced personnel (ANZ, AK) from the NOAA Marine Mammal Laboratory. All deployments were performed with a modified version of the Air Rocket Transmitting System (ARTS, Heide-Jørgensen *et al*. 2001), with compressed air pressures ranging from 9 to 11 bars. The tagger attempted to target a position on the whale that was high on the back, either slightly cranial, or just ventral to the dorsal fin (Fig. 1). A tag carrier separated from the tag on impact and was collected from the water after the whale moved out of the area (Gales *et al*. 2009a).

**Fig. 1.**
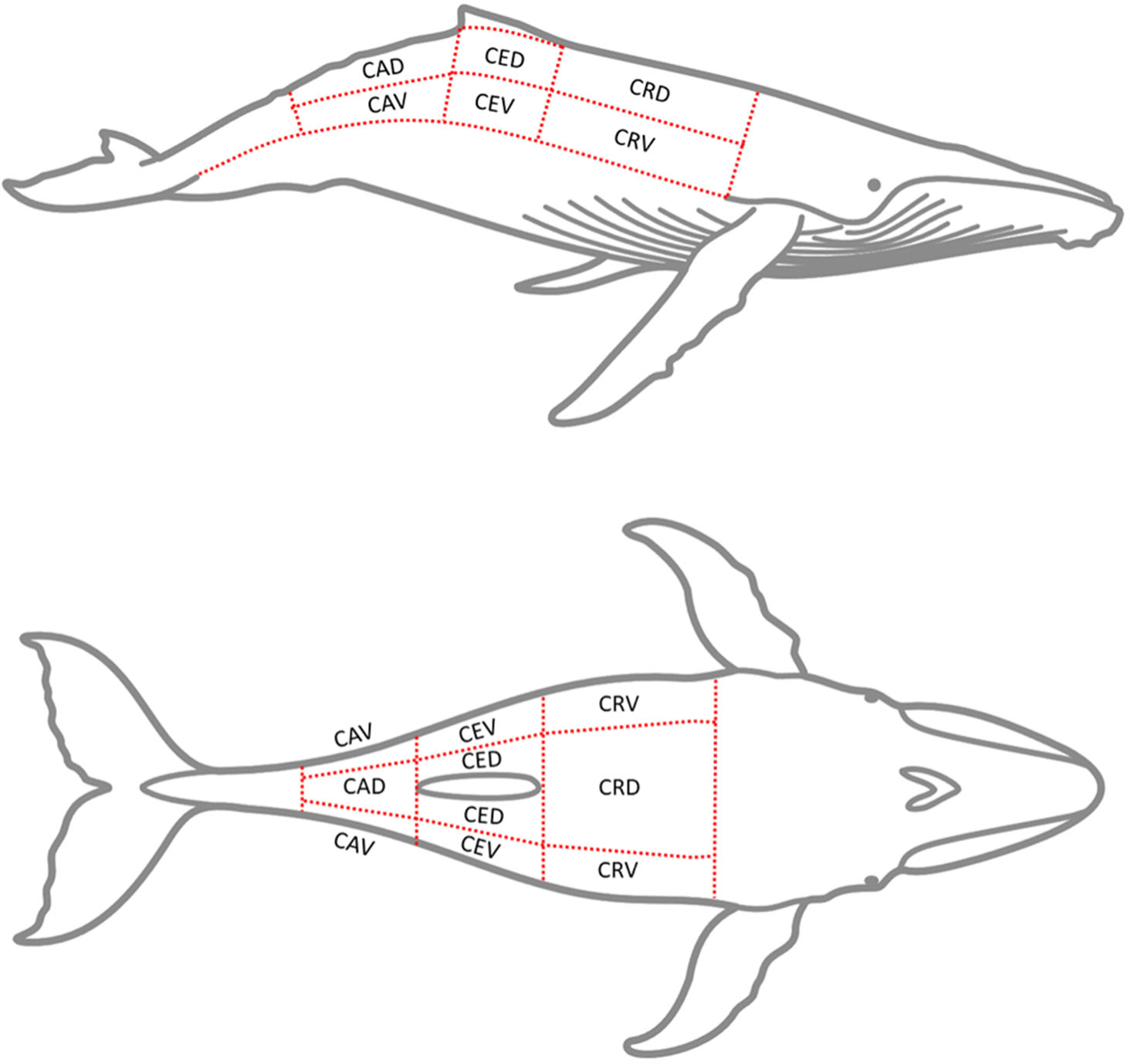
Tag placement by craniocaudal position (CR = cranial, CE = central, CA = caudal) and circumferential position (D = dorsal, V = ventral).

In total, 80 tags were deployed on 79 individuals. One whale was retagged four days after the first deployment because the first tag did not penetrate and was shed immediately after deployment. Whales were photographed prior to, during and after tagging. Focal follows were performed for at least one hour after tagging to assess behavioural responses, injuries and tag placement. Follow-up observations were then attempted on a weekly or bi-weekly basis through December of the tagging year, and then opportunistically in all subsequent years. Follow-up monitoring was also facilitated by a collaborating network of opportunistic observers and commercial whale watching vessels. Images obtained from follow-up were shared with the project team as they became available. The whale identification, tag number, photographed features, photograph date and time, sighting data, and other information such as sampling details were assigned to each image to facilitate image retrieval for analysis. In one case (Tag 1, on day 366), a punch biopsy of skin and shallow blubber close to the tag site was obtained by remote biopsy (Palsbøll *et al*. 1991) and fixed in 10% formalin for histological examination.

### Photograph review

Photographs were used to assess the appearance of the tag wound as well as potentially relevant tag characteristics, such as placement, penetration depth, and retention. These assessments were based on the best available images of each individual per day, hereafter referred to as a “sighting event”.

#### Wound characteristics

Photographs were reviewed by one veterinarian (FMDG), and only photographs of adequate quality (based on focus, resolution, glare, angle) were used to evaluate tag sites. The location on the body, angle and penetration depth were also determined. In each photograph, a series of features were evaluated (skin loss, swelling, exudate, blubber extrusion, depression, colour change, and cyamid presence) and numerical scores were assigned to each feature based on its appearance as listed in Table 1, with higher scores suggestive of more marked changes as illustrated in Figs. 2 and 3.

**Fig. 2.**
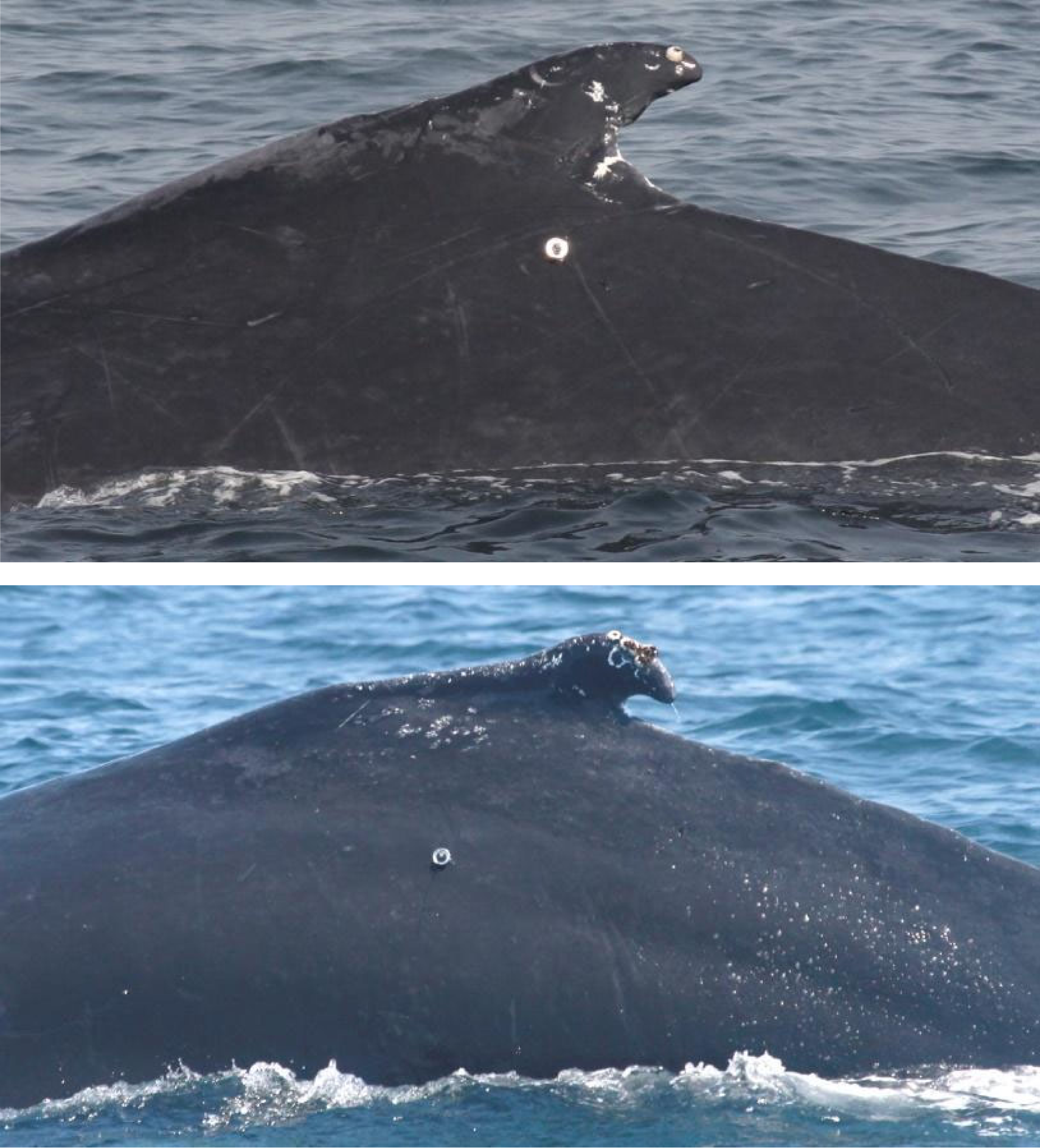
Two GoM humpback whales before any site effects were detected (i.e., Score 0). The cranio-caudal positioning of both was central (under the dorsal fin), but the top image shows an upper flank deployment while the other tag was on the lower flank. Image credits: CCS.

**Fig. 3.**
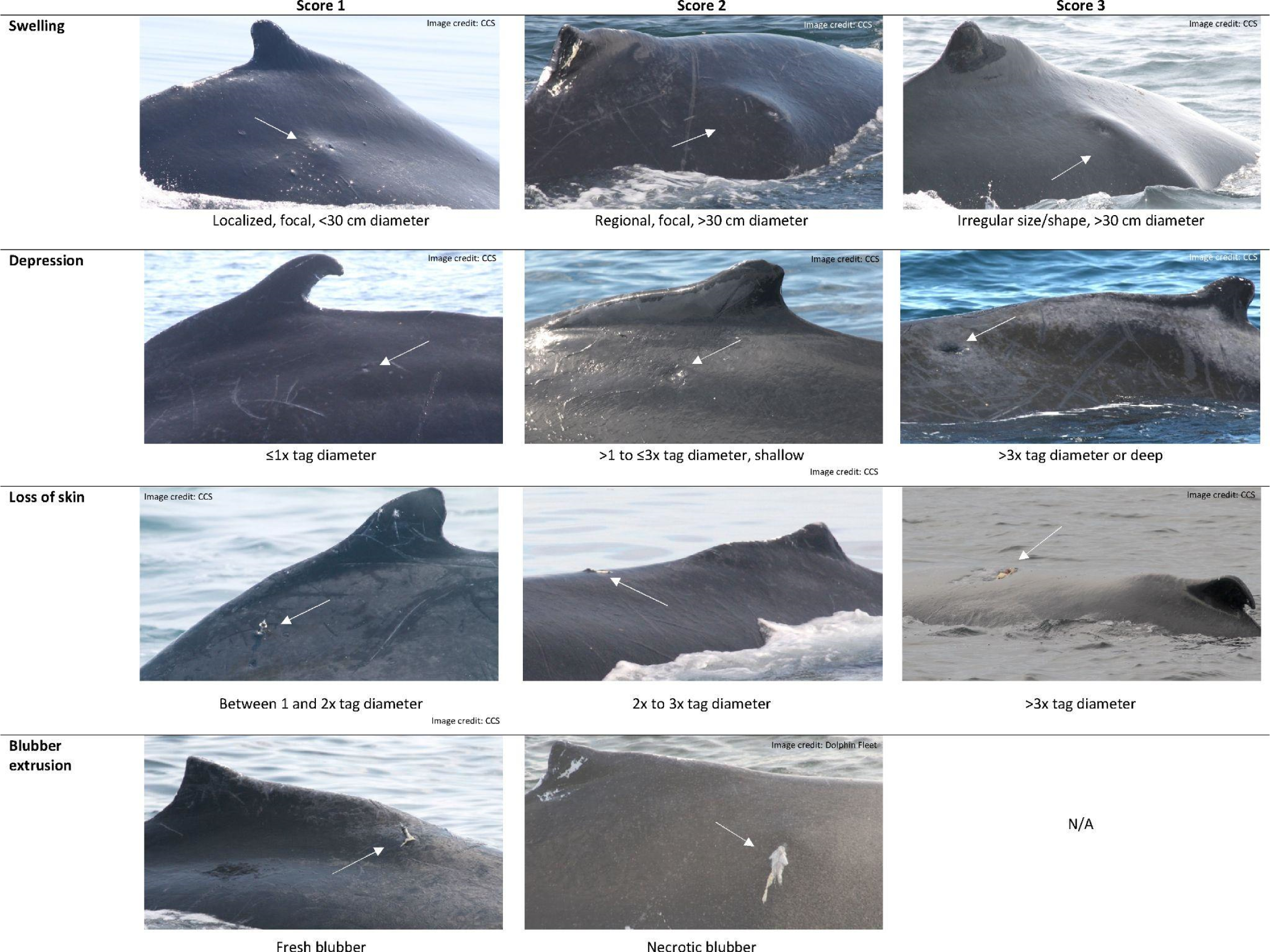
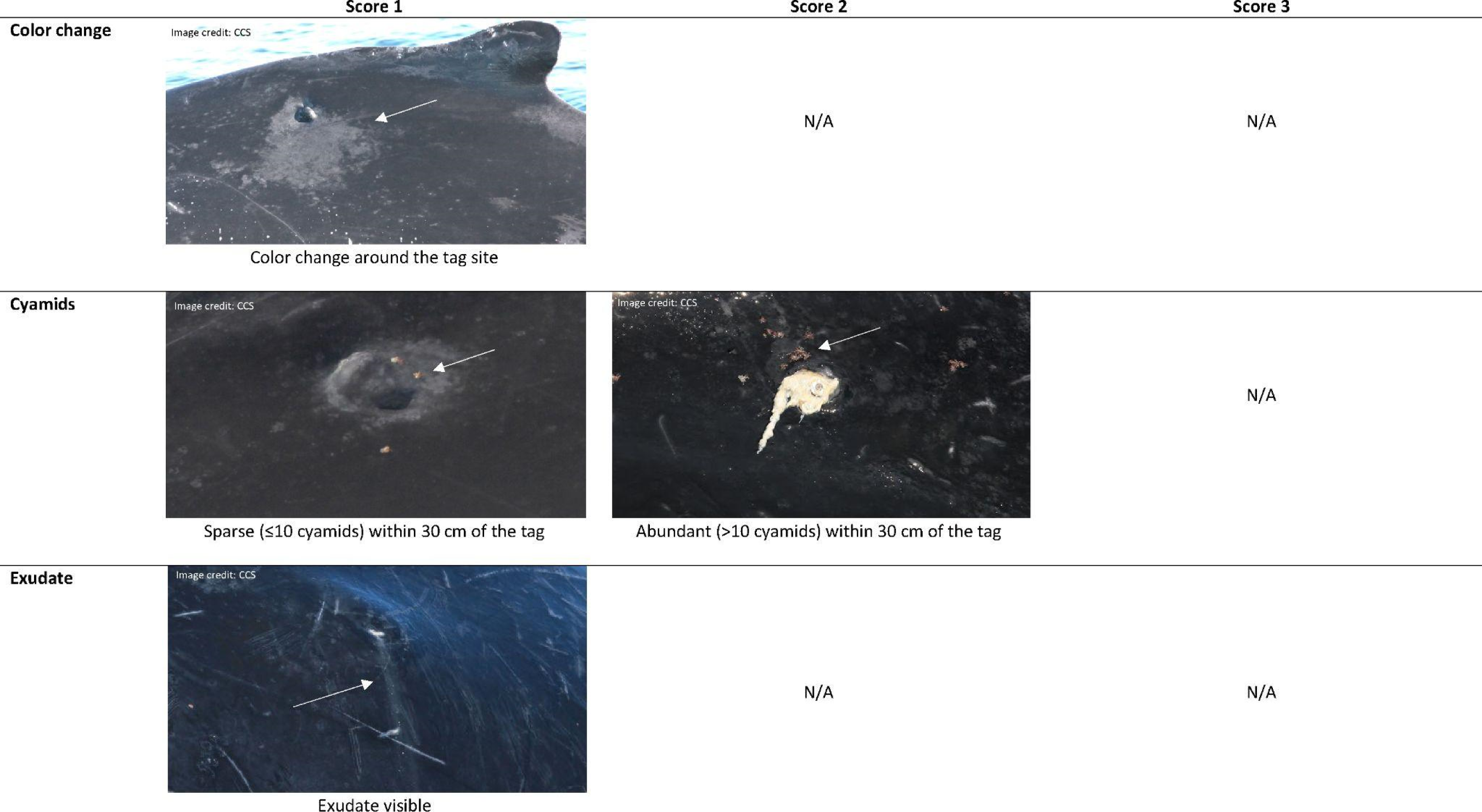
Examples of tag site scoring for all injury features associated with deployment of consolidated satellite transmitters in GoM humpback whales

**Table 1.**
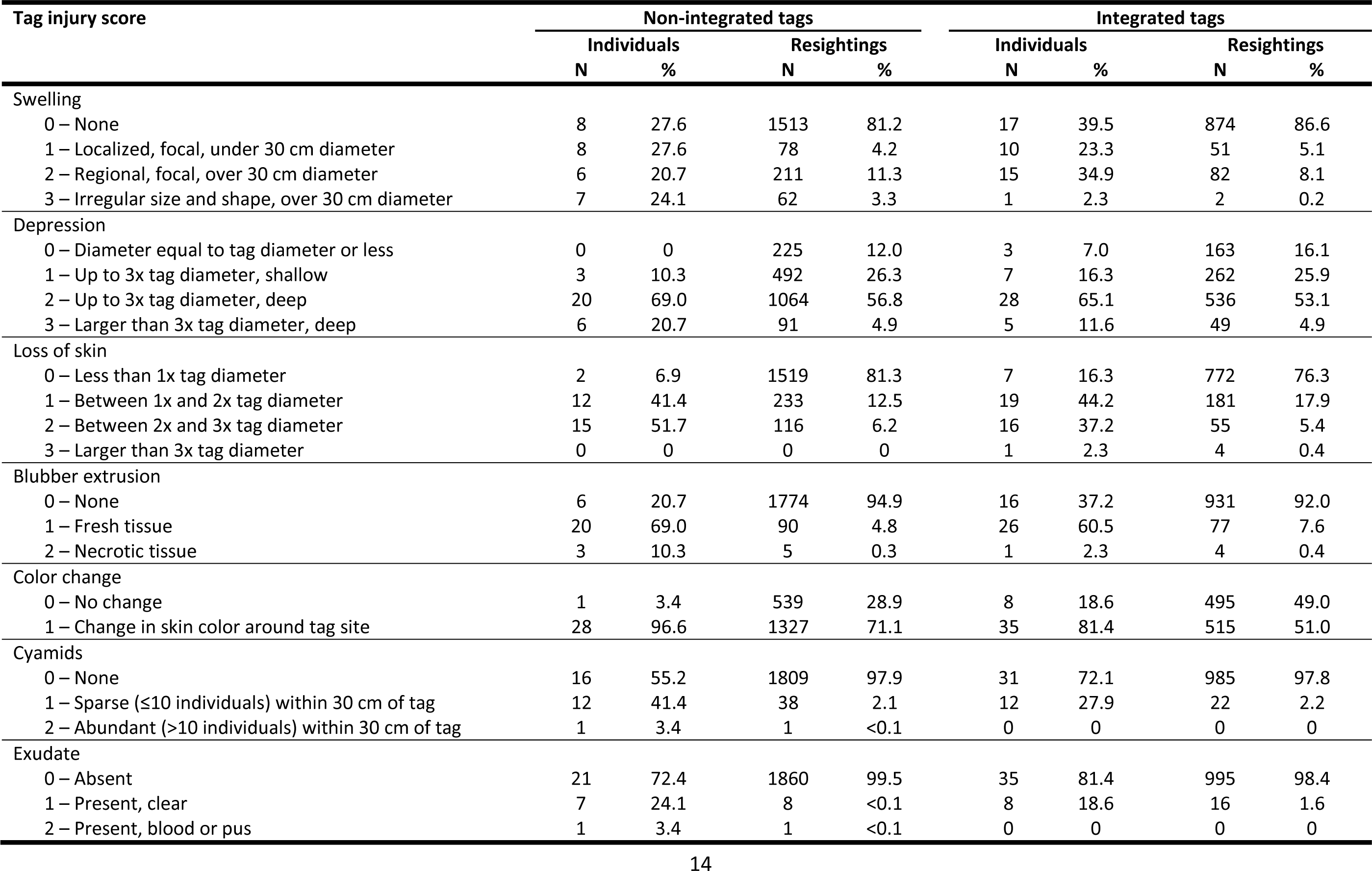
Summary of tag injury definitions and sample sizes for statistical analyses. The latter included scores recorded at the individual level (highest score across all re-sighting events) and for all sighting events. Skin loss scores 2 and 3 were combined in statistical analysis and exudate (presence/absence) was scored but not included in the analysis.

Features of tag sites in different photographs taken on a single day could vary in appearance due to differences in whale body flexion, lighting, presence of water droplets or photographic angle. If a feature could not be consistently evaluated in a photograph on repeat evaluations, it was not given a numerical score, but coded as a “no score”. A multiplier was used to weight total scores on a specific day depending upon whether the lesion appeared to be resolving (multiplier score 0.5, thus the total score was halved) or deteriorating (multiplier score 2, thus total score doubled) compared to photographs from immediately preceding re-sight dates. This subjective value adjustment was useful in highlighting periods of wound change and facilitated the second review of the scores. All photographs taken from 2011 to 2018 were reviewed a second time as a batch in 2019, and special care taken to thoroughly re-evaluate photographs taken during periods of lesion change.

#### Tag and deployment characteristics

Transmission duration was measured as the period between tag deployment and the last transmission received from the tagged whale while tag duration was the maximum number of days of confirmed tag presence. However, a tag part could still be present after transmission ended if breakage occurred. Tag presence was therefore scored as positive when any tag part was visible in any given observation or at any later time. We also assessed the circumferential and cranio-caudal placement on the body. Circumferentially, the tag was categorised as being either in the dorsal fin itself, in the upper (dorsal) flank or in the lower (ventral) flank based on its location relative to a crease that was sometimes visible along the flank and/or an inflection point on the slope of the flank as viewed from behind. The slope was typically flat or slightly concave in the upper area ventral to the dorsal fin and became convex on the lower flank. Cranio-caudal positioning was characterised as cranial (cranial of the dorsal fin and the dorsal hump), central (ventral to the dorsal fin and hump), or caudal (caudal to the dorsal fin). Finally, a photogrammetric method was used to estimate the depth that each tag penetrated at the time of deployment. This estimate was based on the length of tag visible outside the skin, the angle of the tag relative to the plane of the body and the known dimensions of the tag, as described in detail in the Supplementary Material S1.

### Statistical analyses

Cumulative Link Mixed Models (CLMM) were used to evaluate which variables were predictive of each of the six tag injury scores (swelling, depression, skin loss, blubber extrusion, cyamids). Binomial generalised linear mixed models (GLMM) were used for the same purpose for binomial injury scores (colour change). Independent variables considered for this model included sex, timing of the sighting event (elapsed days since tagging), tag deployment characteristics (design, circumferential position, craniocaudal position; Fig. S1) and tag presence upon re-sighting. The squared value of the timing of the sighting event was also included in the initial model to account for potential quadratic relationships. The identity of each whale was included in all models as a random effect. The backward elimination procedure informed by the Akaike Information Criterion (AIC) was used for model selection. Significance level was 0.05 for all tests.

Statistical analyses were conducted using *R* 4.3.1 (R Core Team 2012) with the packages *cowplot* 1.1.1 (Wilke 2020), *effects* 4.2-2 (Fox *et al*. 2022), *ggplot2* 3.4.3 (Wickham 2016), *lme4* 1.1-34 (Bates *et al*. 2015), *MASS* 7.3-60 (Ripley *et al*. 2022), *ordinal* 2022.11-16 (Christensen 2022), and *plyr* 1.8.8 (Wickham 2022). Mann-Whitney and Kruskal-Wallis tests were respectively used to compare maximum number of elapsed days at re-sighting, number of sightings, and tag attachment duration between tag designs and tag models. Local regression (locally estimated scatterplot smoothing – LOESS) was used to summarise the chronological post-deployment progression of tag injury feature scores.

## RESULTS

Of the 79 whales that were tagged, 72 were considered suitable for inclusion in the statistical analysis of tag site effects. Seven whales were excluded from statistical analysis because they had the following atypical characteristics: (a) three whales were tagged on the dorsal fin, (b) one whale had a caudal tag deployment, (c) one whale had a tag that did not penetrate, (d) one whale was never re-sighted after the day of tag deployment, and (e) one whale was re-sighted only twice after tag deployment. The other whales were photographed on a cumulative total of 2,971 sighting events, with an average of 41.3 ± 44.3 sighting events per whale (range = 3 – 218). Sighting events spanned an average of 1156.1 ± 986.4 days (range = 1 – 4110). Tables 1 and S1 (supplementary materials) provide an overview of tag site assessment sample sizes for key whale features, tag design and deployment characteristics. Excluding the dorsal fin deployments, mean depth of penetration was estimated at 238.4 ± 40.8 mm (range = 133 – 294 mm) and 195.0 ± 57.4 mm (range = 42 – 280 mm) for non-integrated and integrated tag designs, respectively.

### Risk factors for tissue site reactions

#### Statistical Analysis

Location and threshold coefficients of the most supported CLLM/GLMM models used to assess severity of the six tag injury categories are summarised in Table 2 and illustrated in Figs 4-9. Model results suggest that the use of integrated tag designs significantly reduces the probability of observing more severe swelling, skin loss and changes in coloration (Table 2, Figs. 4A, 6A, and 8A). Tag design did not influence the severity of depression, blubber extrusion and presence of cyamids (Table 2). Fig. 10 illustrates differences in the predicted score of each tag type across the six tag injury categories over two periods for comparison: the first three years after deployment and the whole study period. Predicted scores show that the deployment of integrated tags is less impactful than that of tags with interfaces or articulations that could lead to breakages.

**Fig. 4.**
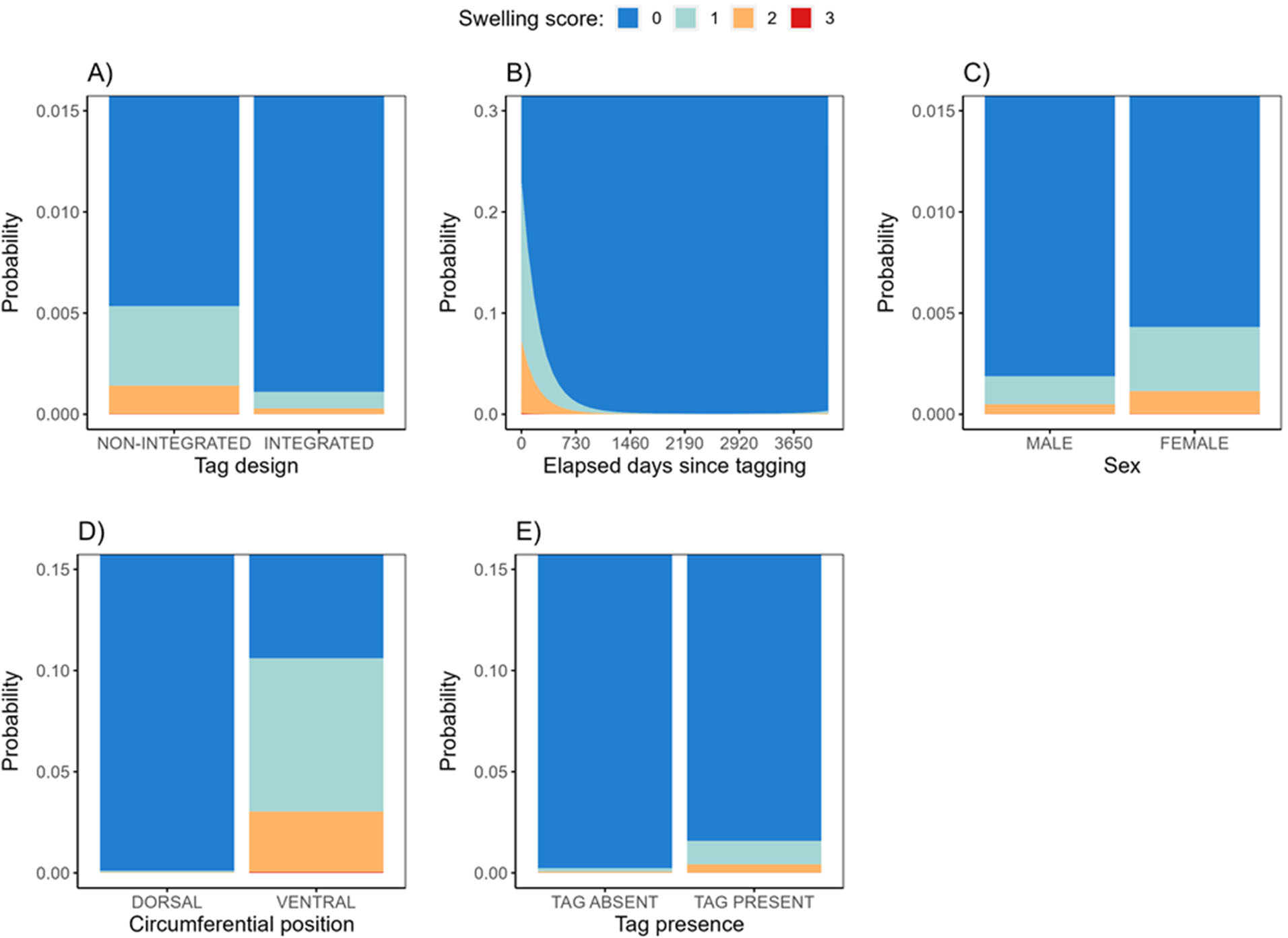
CLMM effects plots for the “swelling” tag injury score: A) Tag design, B) Elapsed days since tagging, C) Sex (not statistically significant, see Table 2), D) Tag circumferential position, and E) Tag presence.

**Fig. 5.**
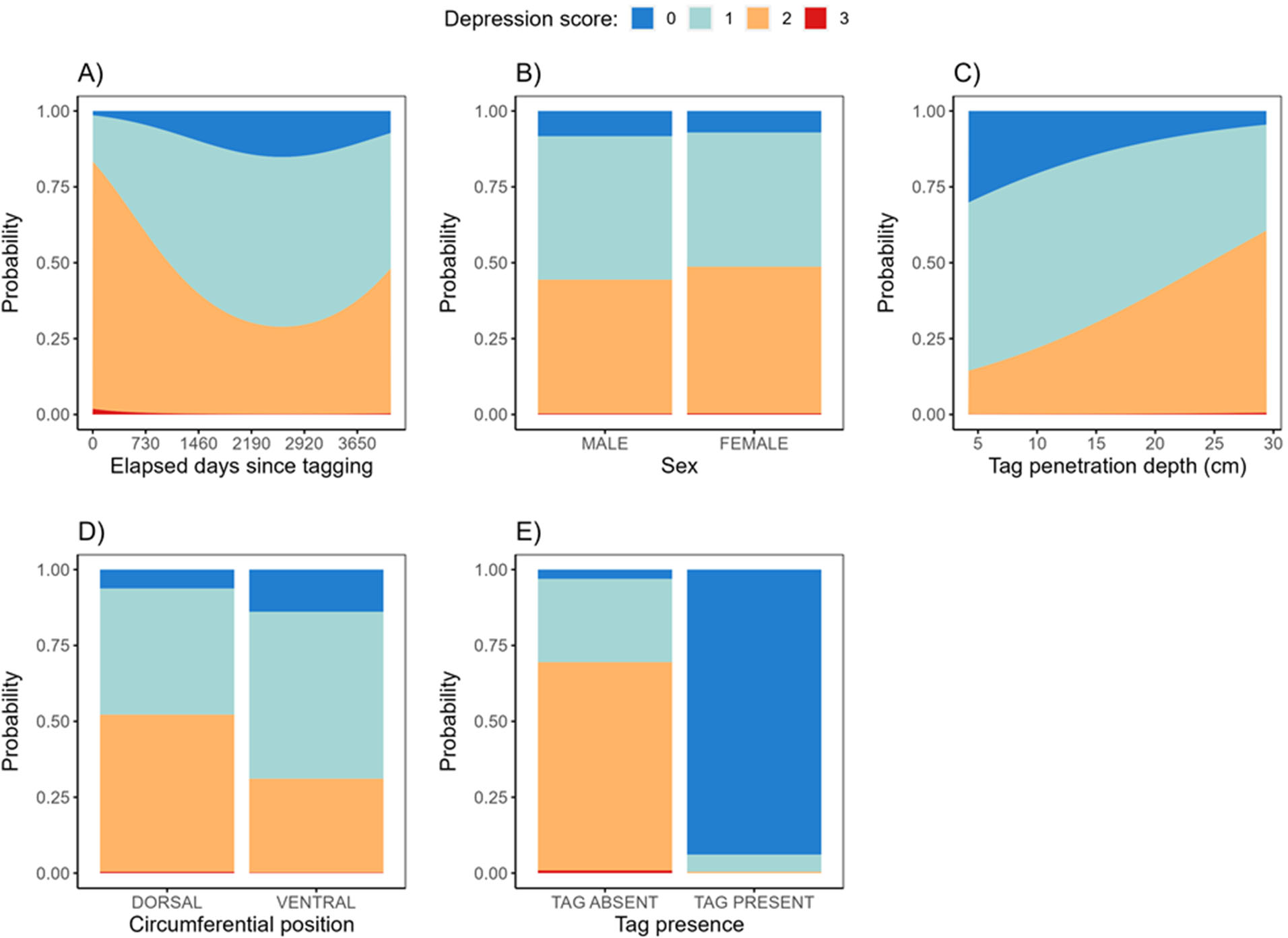
CLMM effects plots for the “depression” tag injury score: A) Elapsed days since tagging, B) Sex, C) Tag penetration depth, D) Tag circumferential position, and E) Tag presence. Only “Elapsed days since tagging” and “Tag presence” were statistically significant (see Table 2).

**Fig. 6.**
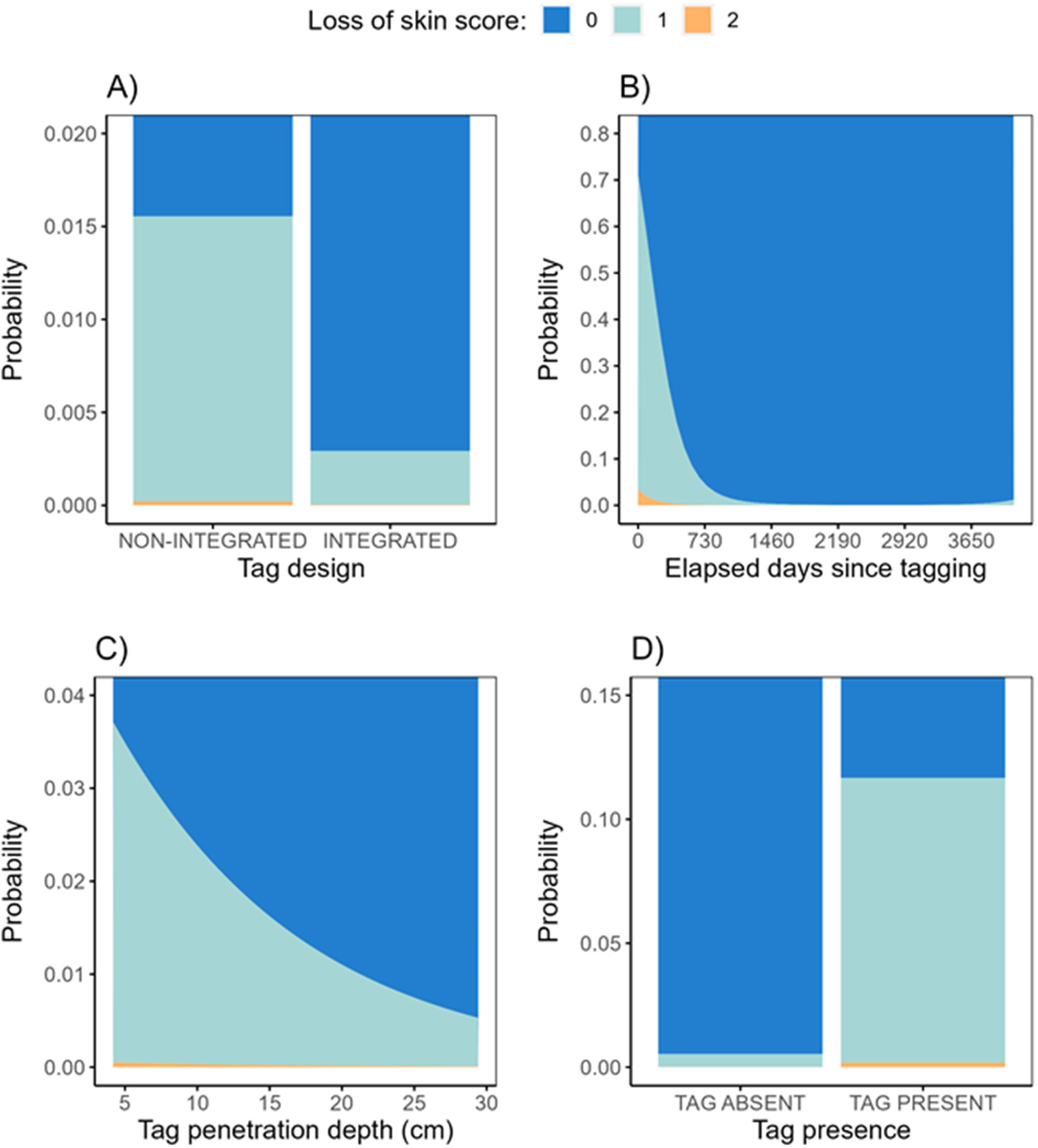
CLMM effects plots for the “loss of skin” tag injury score: A) Tag design, B) Elapsed days since tagging, C) Tag penetration depth (not statistically significant, see Table 2), and D) Tag presence.

**Fig. 7.**
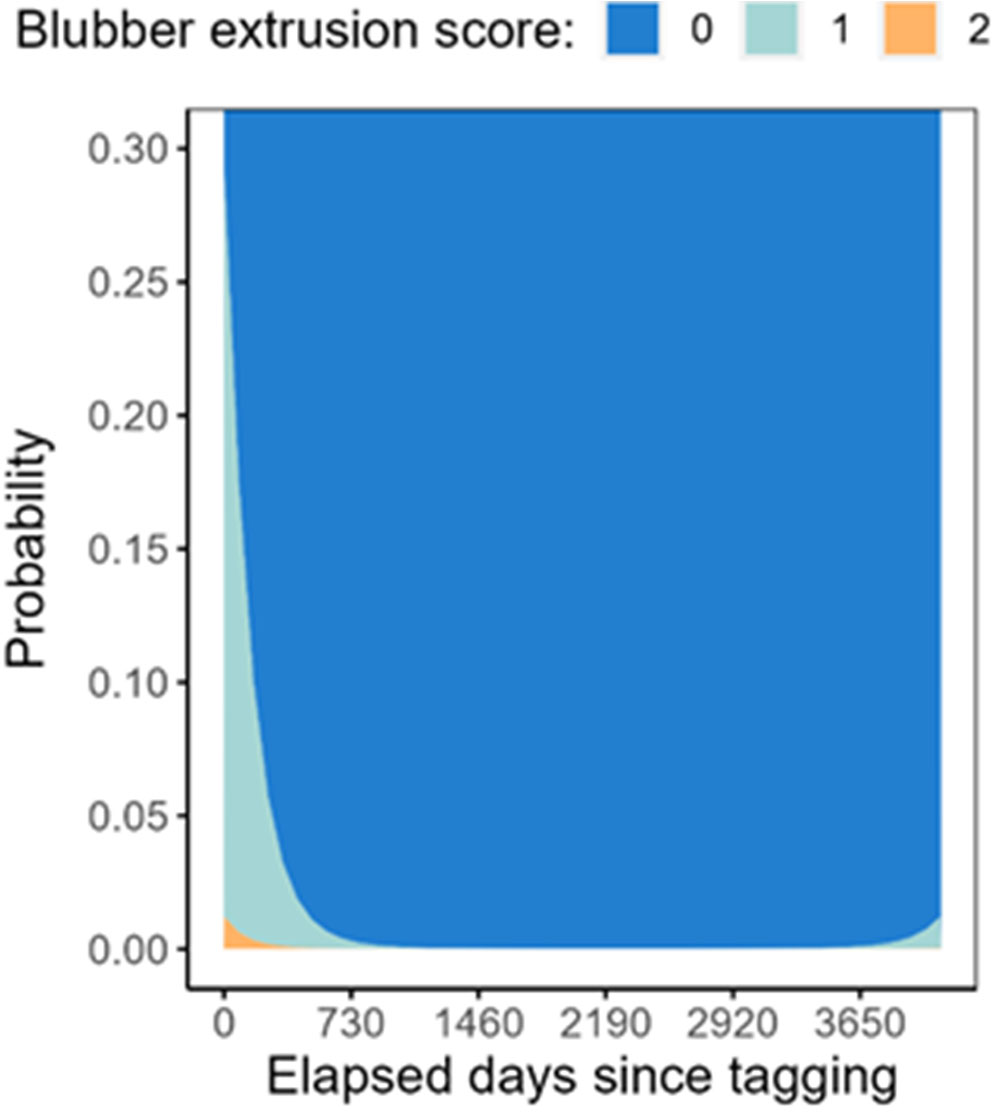
CLMM effect plot of elapsed days since tagging for the “blubber extrusion” tag injury score.

**Fig. 8.**
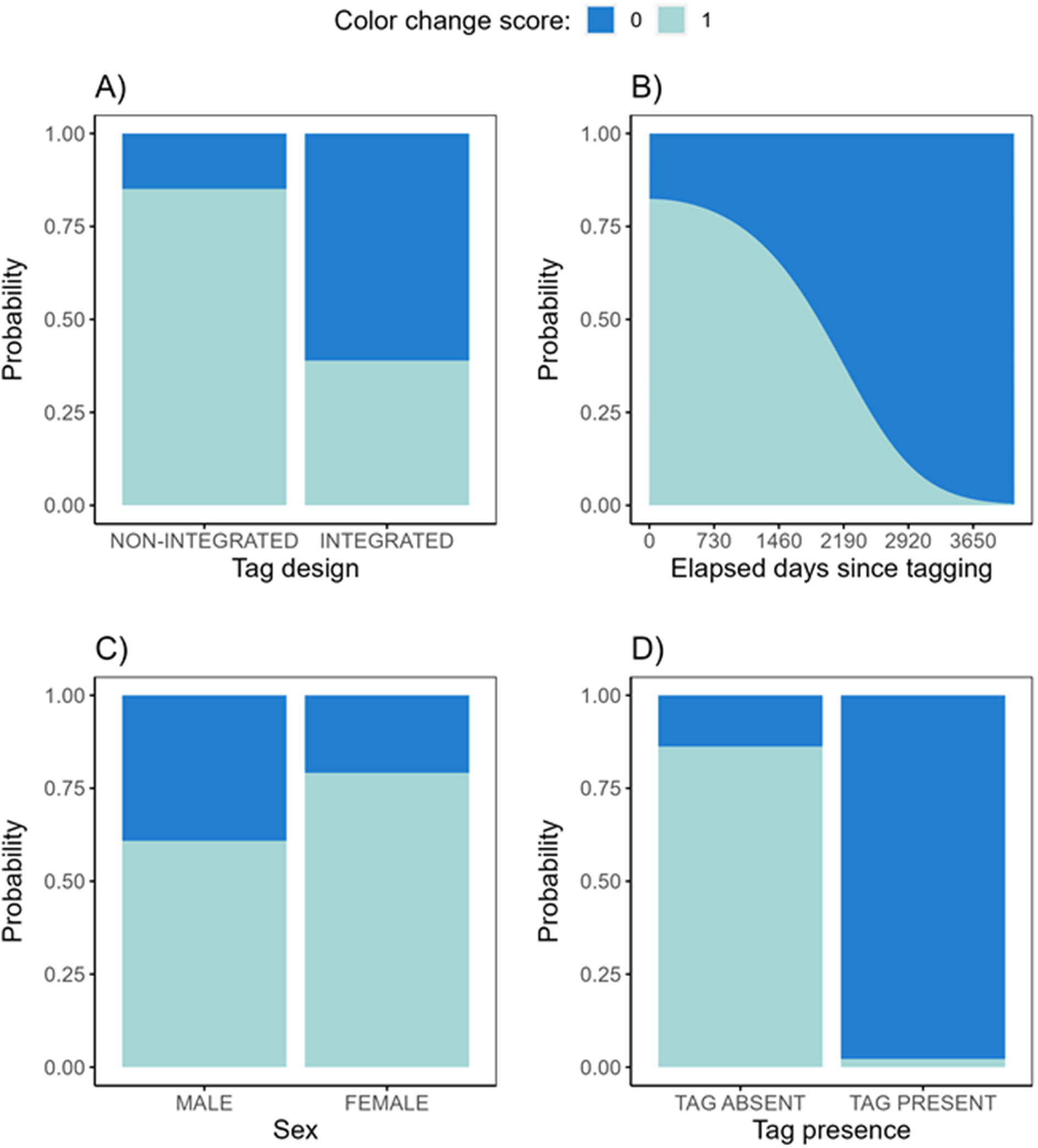
GLMM effects plots for the “colour change” tag injury score: A) Tag design, B) Elapsed days since tagging, C) Sex (not statistically significant, see Table 2), D) Tag penetration depth, and E) Tag presence.

**Fig. 9.**
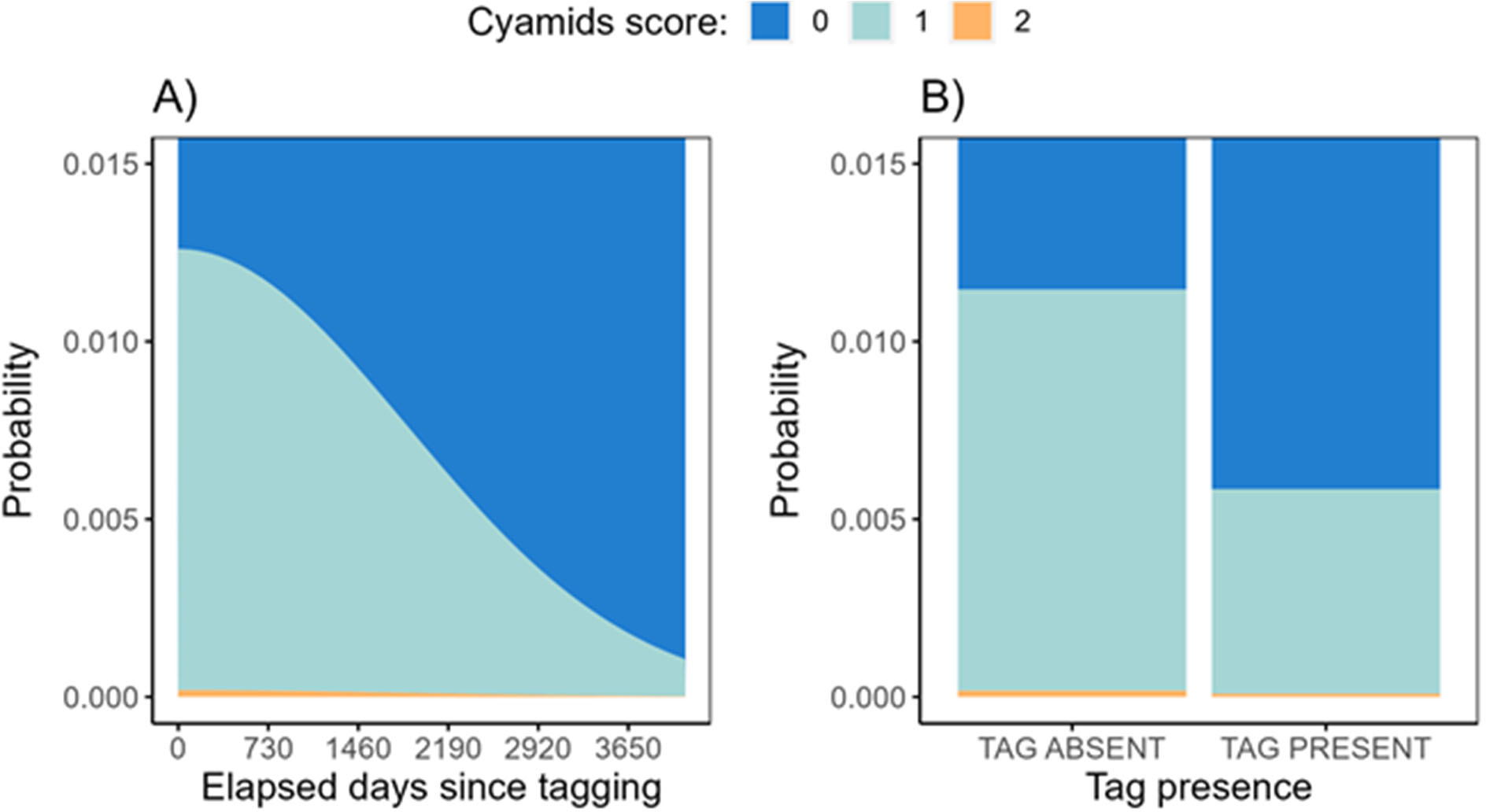
CLMM effects plots for the “cyamids” tag injury score: A) Elapsed days since tagging, and B) Tag presence (not statistically significant, see Table 2).

**Fig. 10.**
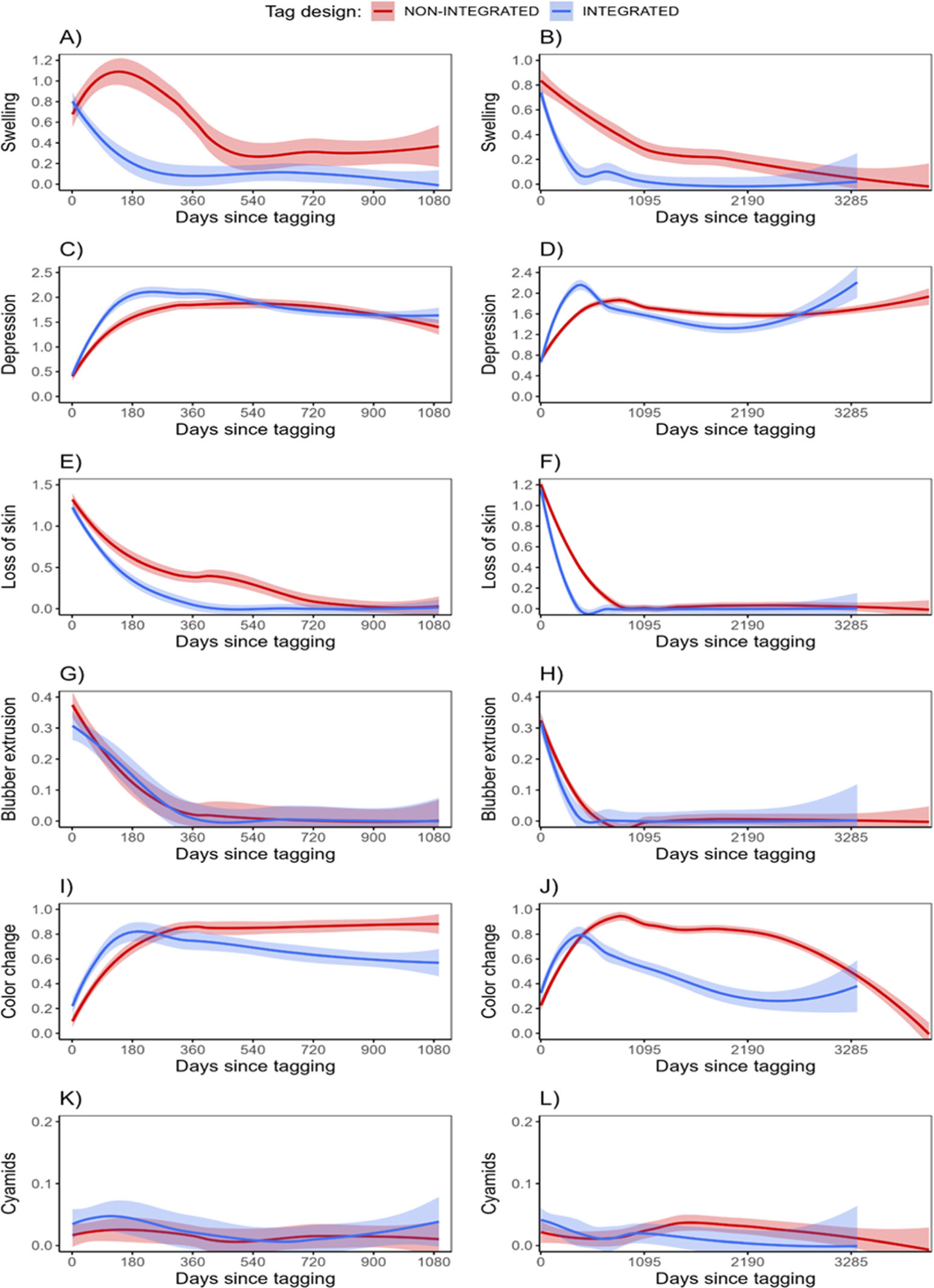
Comparison of the chronology (local regression curve) of tag injury scores among whales that received tags with integrated and non-integrated designs. Lines represent the local regression using data from sightings during the first three years post-deployment (A, C, E, G, I, and K) or during the entire study period (B, D, F, H, J, and L), and shaded areas represent their corresponding 95% confidence intervals.

**Table 2.** Summary of the statistical models to predict tag injury feature scores recorded upon a resighting event. Location and threshold coefficients are expressed as “estimate (standard error)”.

|  | Swelling | Depression | Loss of skin | Blubber extrusion | Colour change | Cyamids |
| --- | --- | --- | --- | --- | --- | --- |
| Model method | CLMM | CLMM | CLMM | CLMM | GLMM | CLMM |
| **Location coefficients** |  |  |  |  |  |  |
| Integrated tag (ref: non-integrated) | −1.580 (0.888) * | . | −1.685 (0.572) ** | . | −2.191 (0.702) ** | . |
| Time since tagging squared (10<sup>3</sup> days) | 0.967 (0.114) *** | 0.368 (0.045) *** | 1.208 (0.113) *** | 1.753 (0.209) *** | −0.426 (0.025) *** | −0.147 (0.064) * |
| Time since tagging (10<sup>3</sup> days) | −5.060 (0.429) *** | −1.925 (0.174) *** | −6.266 (0.431) *** | −8.060 (0.752) *** | . | . |
| Female (ref: male) | 0.834 (0.864) <sup>ns</sup> | 0.173 (0.494) <sup>ns</sup> | . | . | 0.893 (0.689) <sup>ns</sup> | . |
| Tag penetration depth (mm) | . | 0.088 (0.471) <sup>ns</sup> | −0.078 (0.051) <sup>ns</sup> | . | . | . |
| Ventral position (ref: dorsal) | 4.766 (0.958) *** | −0.884 (0.543) <sup>ns</sup> | . | . | . | . |
| Tag present (ref: absent) | 1.948 (0.285) *** | −6.181 (0.226) *** | 3.205 (0.273) *** | . | −5.633 (0.279) *** | −0.679 (0.404) <sup>ns</sup> |
| **Threshold coefficients** |  |  |  |  |  |  |
| Scores 0\|1 | 2.569 (0.913) | −3.259 (1.085) | −2.817 (1.285) | 0.880 (0.178) | −2.640 (0.633) | 4.259 (0.328) |
| Scores 1\|2 | 3.901 (0.921) | −0.639 (1.084) | 1.459 (1.285) | 4.388 (0.376) | not applicable | 8.443 (1.046) |
| Scores 2\|3 | 8.328 (0.962) | 4.925 (1.093) | not applicable | not applicable | not applicable | not applicable |
| **Standard deviation of random effects** | 3.269 | 1.992 | 1.963 | 0.973 | 2.741 | 1.263 |
| **Observations** | 2870 | 2879 | 2880 | 2881 | 2873 | 2855 |

Legend: “.” not included in model, <sup>ns</sup> not significant but included in model, \* P<0.05, \*\* P<0.01, \*\*\* P<0.001.

The severity of all tag injuries was influenced by the time since tagging. Most of the six injury categories were more pronounced at or near the time of tagging, but significantly reduced over time (Table 2, Figs 4B, 5A, 6B, 7A, 8B, 9A). The presence of a significant quadratic term (time since tagging squared) in the models for all tag injury categories suggests that improvements in the severity scores happen in a non-linear fashion as illustrated in Fig. 10.

Tag deployment location was an important predictor for swelling, but not for any other injury category. Model results suggest that tags deployed lower on the body (more ventral position) are likely to result in higher swelling scores (Table 2, Fig. 4D and 4E). The presence of the tag or a tag part (in the case non-integrated tag designs) was positively correlated (more severe scores) with swelling and skin loss; it was negatively correlated (less severe scores) with depressions, skin loss and cyamids (Table 2, Figs 4F, 5E, 6D, 8E and 9B).

Some variables included in the CLMM/GLMM models were not significant (e.g., sex and tag penetration depth, Table 2, Figs 4C, 5B and 5C, 6C, 7B, 8C and 8D), but were retained during model selection for some tag injury scores. This suggests that the models with those variables represent a better fit of the data than those without them and are, therefore, more appropriate for making predictions of severity scores for the relevant injury categories.

### Descriptive assessment of tag site tissue responses

Tag site responses followed a common general pattern: the initial tissue response to tagging was minimal, then progressively there was skin loss around the tag, sometimes a degree of swelling, occasional extrusion of blubber, changes in skin colour, local depression formation, tag loss and skin healing over the tag loss site, but with a depression sometimes remaining (see Fig. 10 for a typical sequence of changes). Although included as a feature in the tag site assessment scoring, exudate was rarely observed, suggesting that photographs are an unreliable method to detect exudation if present, probably due to constant washing of the tag site by sea water. For this reason, exudate detection was considered opportunistic and excluded from statistical models. Cyamid infestation within 30 cm of the tag site was scored, but was rarely severe (when observed, typically 1-5 individual cyamids). The relative frequency of each type of tag site response observed is summarised in Table 1.

#### Non-integrated tags

Tissue responses around tags deployed on whales in 2011 and 2012 were influenced by breakage of the tag at the anchor articulation or at the interface between the anchor and transmitter. Tissue reactions were initially minimal, but following transmitter loss, lesions typical of foreign body reactions slowly developed. In 11 cases, tag breakage was visually confirmed, and typically (n=8) involved the retention of a fragment of, or the whole anchor, after transmissions stopped (Figs 11 and 12). Confirmed observations of post-transmission part retention were made up to 846 days (mean=167.7, SD=275.9) post tagging. Two of the eight cases were deployments in the dorsal fin and a third was the only caudally-placed tag. The 24 whales without observed tag fragments had tissue reactions similar to these 8 whales and the sequence of tissue responses suggested tag breakage also occurred in these whales.

**Fig. 11.**
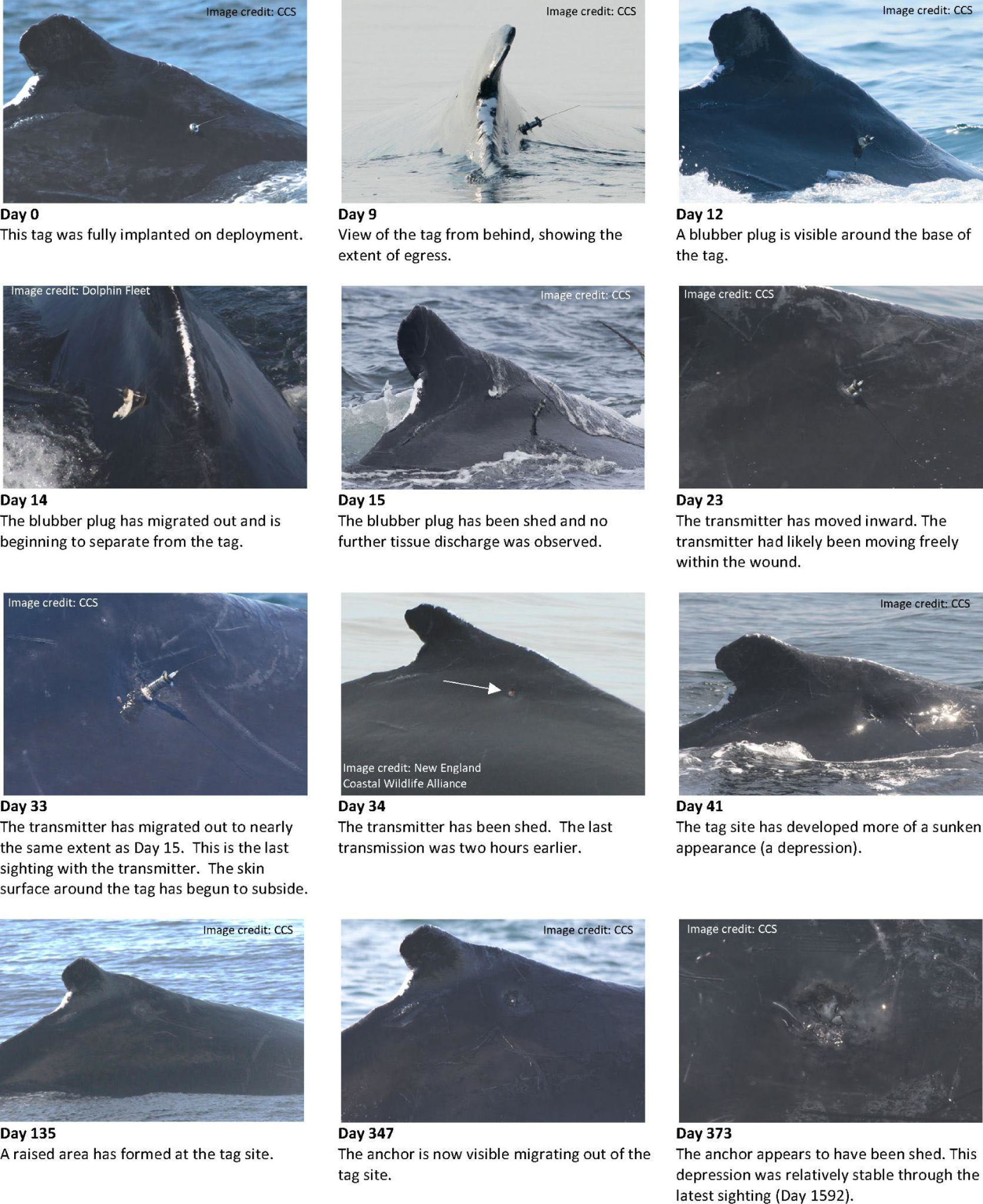
Typical sequence of changes in the tag site during tag rejection and healing in a humpback whale instrumented with a non-integrated satellite transmitter in the GoM (Tag 3).

**Fig. 12.**
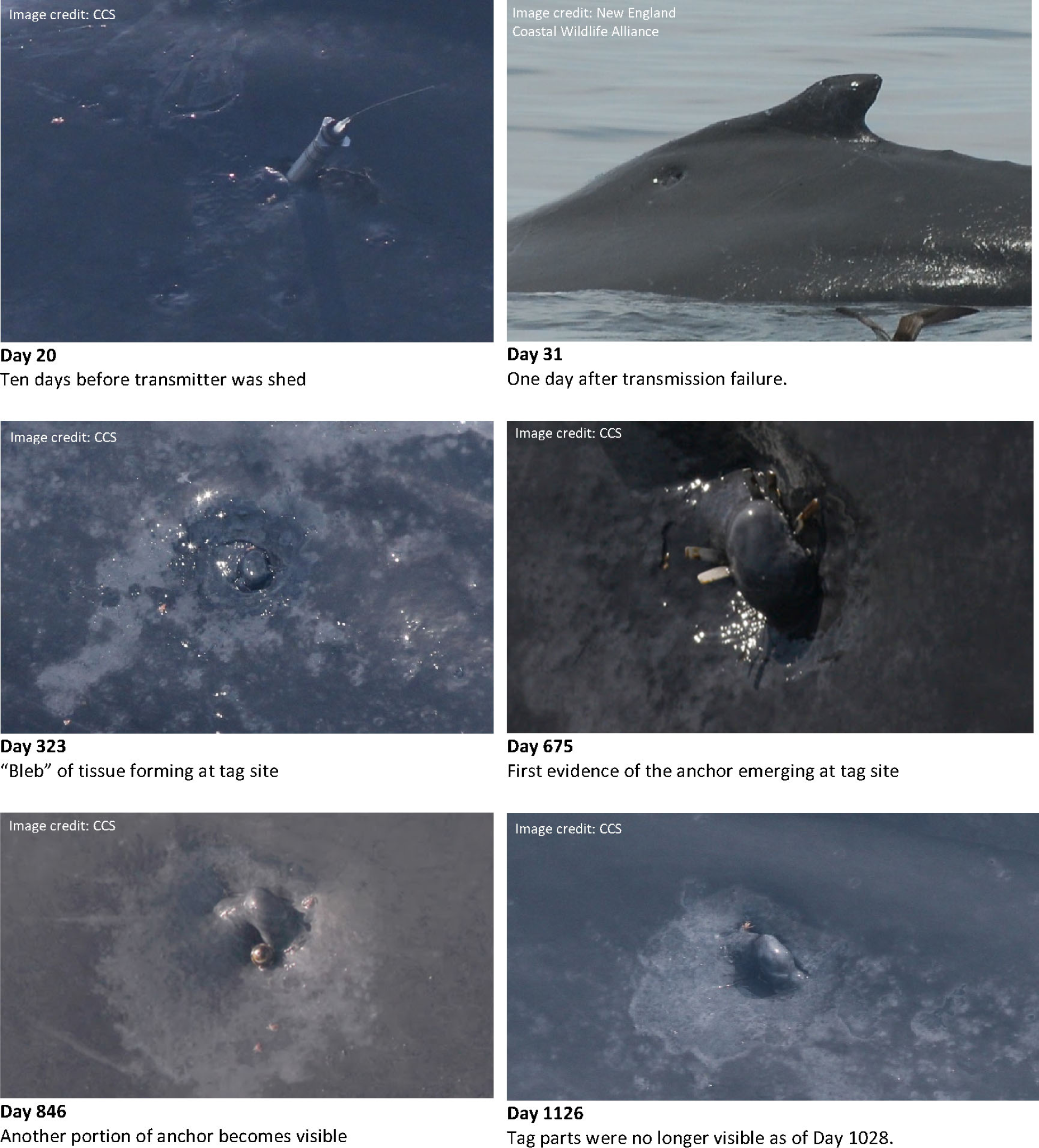
Example of tag site appearance during rejection and healing in the presence of broken tag parts on a humpback whale tagged with a non-integrated satellite transmitter in the GoM (Tag 5).

The progression of tissue change scores following tag insertion are illustrated in Fig. 10. Tag insertion was associated with minimal observable immediate skin reaction and no obvious blood loss, with skin margins adherent to the tag at the insertion site. In four whales, there was slight enlargement of the area of perforation in the skin around the tag after 1-4 weeks, resulting in a skin defect up to two times larger than the diameter of the tag that lasted several months. In most cases, however, there was minimal enlargement of the hole in the skin around the inserted tag, with skin edges abutting tightly against the body of the tag, suggesting minimal movement of the section of tag perforating the skin. In three animals, at about a year (and in one animal, Tag 10, at three years), the skin defect around the tag enlarged to about three times the tag diameter.

Swelling around the tag occurred in eight tagged whales within days of insertion. These swellings developed rapidly over 1-3 days, were a uniform shape and size in five animals, irregular in three, and persisted for a month to seven years. When tag remnants were visually confirmed, the sizes of the swellings appeared to remain relatively constant over the years until the remnants were no longer detected. The body condition of the whales, however, influenced the appearance of the swellings, which were more marked in the first half of the feeding season (through July) compared to later. Presumably, the increased blubber thickness around the tag site masked the appearance of the swellings.

In 11 cases, pieces of blubber were observed protruding from the tag wound site as the tag transmitter was shed. Tissue other than blubber, such as muscle or connective tissue, was never observed extruding from the tag site. Protruding blubber was initially white and gradually became grey-yellow, especially at the distal end of the tissue furthest from the body of the whale, suggesting gradual necrosis of the fragment due to loss of blood supply. Once these pieces of blubber were separated from the body, skin margins began to heal over the tag site. The skin surface was depressed at the healed tag site and this was often surrounded by variably shaped patches of discoloured skin. The depression at the tag site was sometimes so deep that it was not possible to determine whether or not there was ulceration of the skin within the depressed area. Cyamids were occasionally present within these depressions. Over months, the depressions became less marked, and were hard to detect by 2022 (see Table S2).

In one-third of the 35 whales, small, raised area, or bleb, of smooth skin were observed within the depressed area at the tag site one to three years after tagging. In some cases, the blebs disappeared over months to years. In other whales, they persisted and were observed 10 years after tag application. In one tagged whale for which frequent sequential photographs of tag fragment rejection were available (Tag 05), the fragments did not extrude from the bleb of skin, but adjacent to it (Fig. 12). In a whale that was tagged on the dorsal ridge (Tag 17), the tag appeared to have been extruded from the contralateral side from application and a bleb was observed at the exit site.

The first whale tagged in 2011 exhibited a strong, extended behavioural reaction to tagging, and although the tag had fully penetrated, the transmitter was shed by the following day. The exact reason for this outcome is not known with certainty, but it could have been due to tag breakage. No tag parts were ever observed in photographs, but the whale’s nutritional condition declined over two months post tagging, and a persistent, regional (score 2) swelling developed surrounding the tag wound site. A purulent exudate was observed once, 366 days post tagging. Histology of a biopsy sample collected near the tag site revealed mild dermal fibrosis, perivascular edema and minimal neutrophilic dermatitis. Special stains (periodic acid-Schiff and Gomori’s methylene silver) for fungi were negative. The last assessment of this whale was 391 days post tagging, when body condition had improved since earlier that summer, and the whale has not been seen since then. The regional swelling, presence of a purulent exudate and mild dermatitis suggest that infection of the tag site could have occurred.

In five cases, the tag was deployed on the dorsal fin (Tags 02, 23, 30 and 32), or the dorsal ridge (Tag 17, also discussed above). In two of those cases (Tags 02 and 17), discoloured skin was observed on the contralateral side after tag loss, with focal depression of the skin on that side. These changes in skin colour and texture could result from changes in blood supply to the area, migration and exit of a tag fragment to the contralateral side, or contraction of scar tissue on the tagged side pulling tissue internally from the contralateral side affecting skin integrity. By 10 years after tag loss, there were no detectable changes in the skin at the tag site or the contralateral sides compared to surrounding tissue in these five whales.

At last observation, 11.8% (n=4) whales tagged with non-integrated tags had barely detectable marks on the skin, 58.8% (n=20) had small shallow depressions in the skin (1-3 times the diameter of the transmitter body), 26.5% (n=9) had deeper depressions with areas of skin with differing colour from the adjacent skin, and 2.9% (n=1) had a mild swelling around the tag site (Table S2). The remaining whale was observed only once, on the day after tagging, and there was a mild swelling at the tag site at that time.

#### Integrated tags

Forty-five integrated tags were deployed on 44 whales in 2013, 2015 and 2018. The chronology of the changes in features evaluated in each photograph is shown in Fig. 10 and a typical sequence of events illustrated in Fig. 13. Immediately after tag deployment, minimal tissue responses were observed. Rarely, drops of blood were observed in photographs taken within an hour of tagging, but most often, no discharge was observed and the skin margins appeared tight against the transmitter body. Within days of tagging, the skin margins typically appeared separated from the tag with a skin defect of up to twice the diameter of the transmitter body, with no discharge or tissue extrusion observable. Over the following weeks, variably sized swellings were observed around 25 tag deployment sites, while no swelling was observed at 20 tag sites. In nine whales, swellings were Score 1, fifteen whales had transient swellings scored as Score 2, and one whale had a Score 3 swelling. In six whales (Tags 46, 51, 54, 57, 61 and 64), small pieces of blubber were observed around the tag that gradually became discoloured and were extruded 2-4 weeks post tagging. Once the tag was ultimately shed, the skin healed over the tissue deficit, and a depression in the skin surface was apparent at the tag site. Occasionally the skin around the healed tag site was paler than adjacent skin. The size of the depression and area of coloration change corresponded to the sizes of the swellings and areas of skin lost during tag retention. A raised skin bleb within the depression was observed in one whale.

**Fig. 13.**
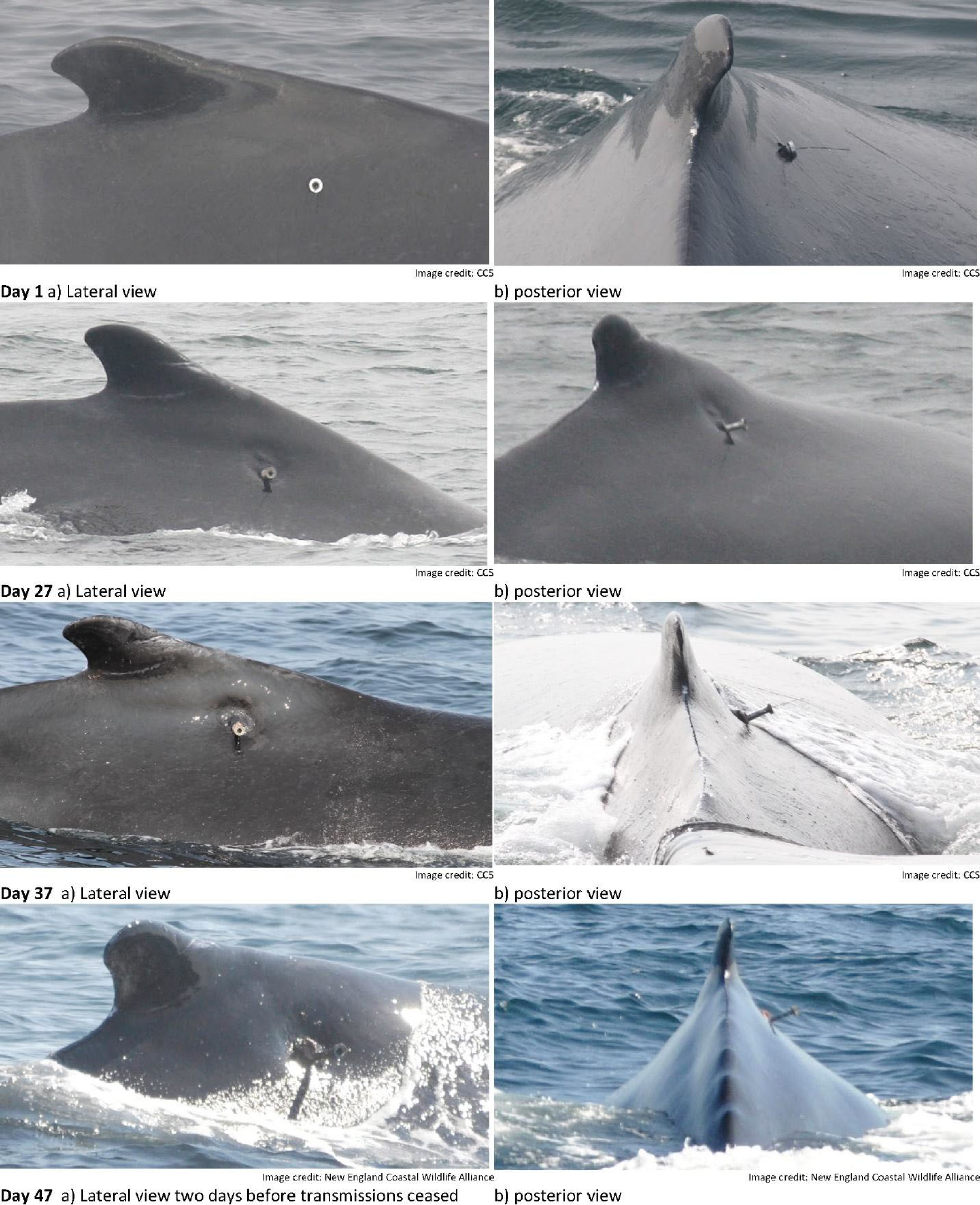
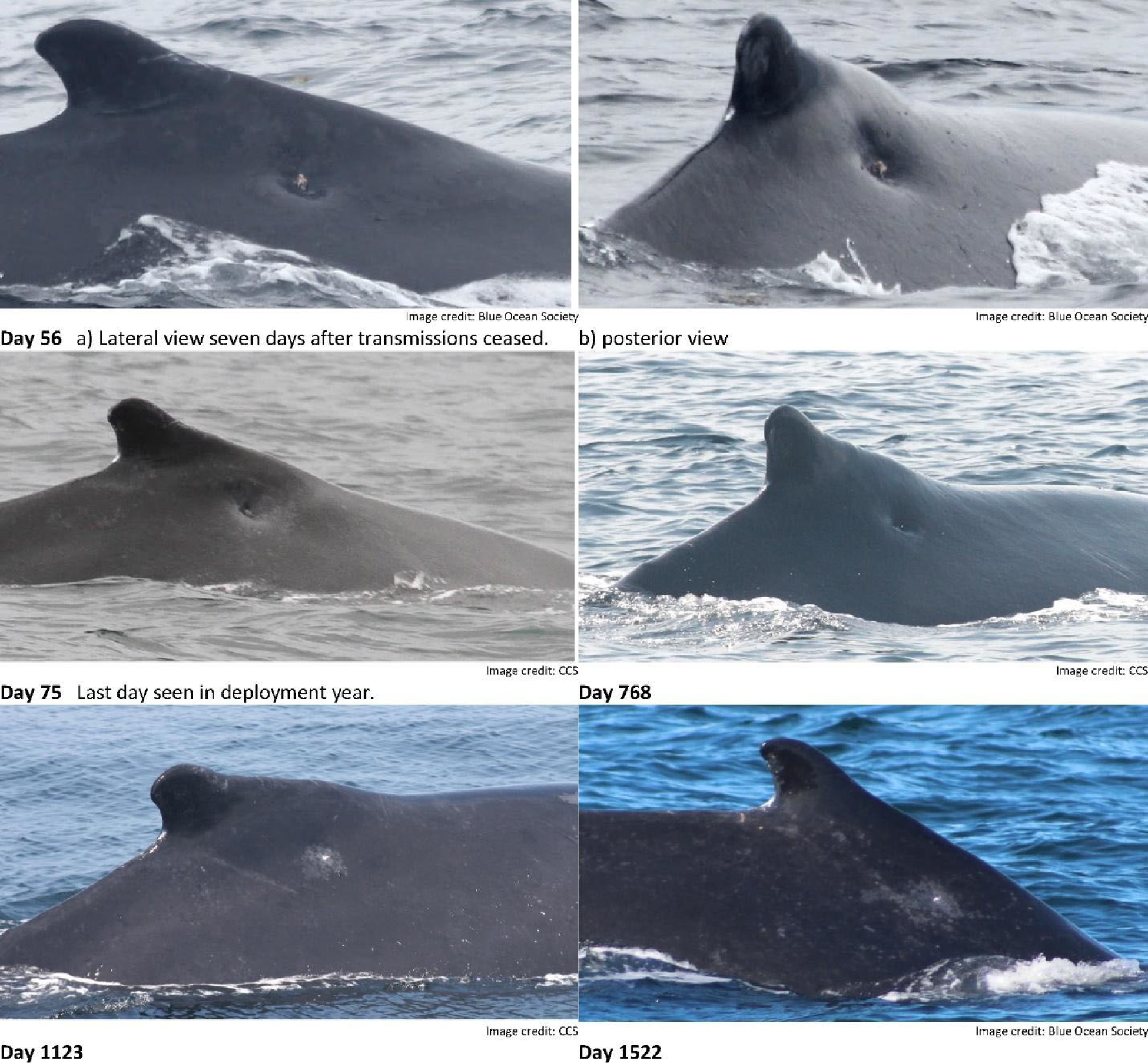
Example of non-integrated tag rejection and healing on a humpback whale tagged with an integrated satellite transmitter in the GoM (Tag 68).

By one year after tag loss, the skin depressions at the tag sites were visible in high quality photographs, but some could be missed in photographs taken at greater distances, or those of poor quality. In one whale in which the tag was applied just right of the dorsal midline (Tag 79), marked depressions were observed on both sides of the dorsal ridge, with the depression in the contralateral side being larger than the one at the site of tag insertion.

One whale (Tag 48) tagged very close to the midline cranial to the dorsal fin had minimal immediate tissue reaction to the tag. At day 56 post tagging the tag was no longer visible and an irregular-shaped wound was present around the tag site that was approximately 50 cm in diameter, with multi-focal irregular shaped areas of skin loss and exposed blubber. This healed over days 57-90, and at day 346 the site had healed leaving an irregular shaped area of rough skin. It is likely that this tag was caught on a foreign object or rubbed out of the animal, pulling tissue with it as well as abrading other tissue, to leave this irregular shaped lesion. No evidence of purulent discharge or foreign body in the wound was detected.

An unusual egress event occurred in the case of one integrated tag (Tag 69). This individual was entangled in a boat anchor line 10 days after tagging in 2018, and the tag was detached by the entangling rope. The tag site was well-documented minutes prior to the event as well as shortly after the tag was removed. Images show rope starting in the mouth and trailing aft along the body, sometimes at the tag site. It is our assumption that the entangling gear dislodged the tag, and there is no evidence that the tag was a direct or indirect cause of the entanglement. Although there was initially bleeding at the tag site following tag loss, no unusual tissue damage was evident at the skin surface. A disentanglement team responded the same day and removed a large portion of the life-threatening gear, but not all of it. This individual was never re-sighted, but was retrospectively matched by CCS to a whale that was found dead from unknown causes in 2019. No necropsy was conducted in that case.

At last assessment of the 45 whales tagged with integrated tags, 15.6% (n = 7) had no obvious mark on the skin, 66.7% (n = 30) had small, shallow depressions (score 1), 15.6% (n = 7) had deeper depressions (score 2) and 2.2% (n = 1) had mild swelling at the tag site (Table S3).

## DISCUSSION

The series of photographs in this study provide the most comprehensive description of tissue changes around satellite tags in baleen whales to date. However, assessments are limited by being based on photographs and typically lacking insights from histological or ultrasound examination of tissues. Thus, interpretation of the significance of features such as swelling is limited to the sequence of their apparent changes over time, potentially further informed by other data, such as whale behaviour, reproduction, body condition and survival. The availability of many photographs of tag sites in individually identified whales over years enabled the development of an objective scoring system to characterise temporal changes at tag sites. This scoring facilitated comparison of tissue responses amongst whales and tag types, so that factors minimising tissue responses could be identified. Although the addition of a multiplier that was based on the decision of the scorer as to whether the lesion was improving or worsening was subjective, it was useful in focussing attention of the reviewer to periods of change when reviewing hundreds of photographs spanning years. In humans, an objective scoring system for evaluating surgical wound healing using only appearance has been useful for evaluating surgical success, and for identifying risk factors for prolonged wound repair (Bailey *et al*. 1992; Gorad *et al*. 2021). In wildlife, such a numerical system has only rarely been used, yet has been proposed as a tool for cross-ocean comparison of wounds in whale sharks (Womersley *et al*. 2021) and could similarly be useful for evaluating impacts of different tag types in cetaceans.

The series of observations in the first years of this study enabled improvements in the tag design to prevent tag breakage, prolonged retention of tag fragments, foreign body reactions and delayed wound healing. Modifications of the tag design through the study resulted in improved skin healing and minimal residual tissue changes in whales associated with the integrated tag design. The effort undertaken to re-sight tagged whales repeatedly was invaluable to improving tag designs and understanding the causes of prolonged tag site healing. We have yet to fully quantify the forces on tags during deployment and within the body. Consequently, we recommend that tag design and manufacturing techniques avoid the use of elements that might break as well as those that could result in tissue shearing.

The tissue responses observed in whales tagged with integrated tags suggest there was moderate local reaction to the insertion of an indwelling integrated tag placed dorsally in the proximity of the dorsal fin of a humpback whale, with wound healing by second intention once the tag was shed. Typical wound healing by second intention (filling a tissue deficit) involves epithelialization, angiogenesis, collagen formation and finally contraction, in contrast to primary intention healing which occurs when skin margins at a wound site are closely aligned by suturing. The discoloration of the whale skin around the tag site weeks after tagging is likely a consequence of new skin cell production (epithelialization) and new blood vessel formation (angiogenesis), while the depressions observed months to years after tag loss likely result from contraction of the wound site following blubber loss and tissue healing. The duration of each phase of wound healing observed after tag loss in this study fit within the expected durations of phases of wound healing observed in large cetaceans injured by other traumatic events (Bruce-Allen & Geraci 1985; Zasloff 2011). Exceptions were the skin blebs observed in association with broken tag fragments that may be epithelial masses similar to keloids observed in people with delayed skin healing (Murray *et al*. 1981).

Potential causes of the swelling around the tag site are bleeding with haematoma formation, tissue edema, inflammation in response to trauma or potentially salt water intrusion, pus accumulation, or a mixture of these. Bleeding or purulent discharge were rarely observed, making hematomas or abscesses unlikely causes of the swellings. Although pus was not observed, bacterial infection around embedded tag parts has been detected by histology of tissue from beluga whales (*Delphinapterus leucas*) tagged with “spider” tags (Burek-Huntington *et al*. 2023). Thus, photographs alone showing lack of purulent discharge are not conclusive evidence of lack of infection, although it is unlikely in these humpback whale cases given the progression of the lesion development. More likely causes of swelling are edema or inflammation due to mechanical trauma to tissue, as any disruption of the keratinocyte layer at the skin surface will initiate an inflammatory reaction (Nickoloff & Naidu 1994). Surgical-grade sterilisation (versus disinfection) of the tags prior to use was likely important in minimising infection post insertion, and is a practice that should be continued despite the additional logistical challenges involved.

The tags used in this study were longer than the minimum distance across the blubber from outer epidermis to underlying muscle, as adult humpback blubber thicknesses at tag deployment sites reportedly vary between 10 and 18 cm seasonally (Slijper 1962; Lockyer 1981; Gabriele *et al*. 2021). There are no recent published data on blubber thickness of adult humpback whales in the GoM, but insights can be drawn from two stranding events involving catalogued adults in recent years. One adult male had a 7 cm blubber layer at the mid-lateral thoracic region in the month of January (AMCS013Mn2013, AMSEAS unpublished data). An adult female ranged from 6– 7 cm blubber thickness across two common tag placement locations in May (IFAW19-287Mn, IFAW/CCS unpublished data). This information, combined with our photogrammetric tag penetration estimates suggest that most tags in this study likely extended below the blubber. The post-mortem examination of a North Atlantic right whale that had been darted with a syringe and needle to administer antibiotics about a week prior to death revealed the potential for a rigid object traversing the blubber-muscle interface to cause tissue shear and cavitation (Moore *et al*. 2013). Further studies on dolphin carcasses indicated insertion of a rigid needle across the blubber-muscle interface can cause cavitation in the muscle due to shearing action, with the effect more marked further from the dorsal midline (Moore & Zerbini 2017). Thus, it is possible that the tags were causing cavitation with associated hematoma formation, the swellings being haematomas in the connective fascial layer. However, the speed of swelling development and the lack of change in size over time makes this less likely than tissue edema following trauma. A 16-year time series of photographs of a female blue whale tagged with an old model Type-C tag had a persistent skin lump associated with a retained tag fragment for a decade, yet later shed the tag and the lump resolved (Gendron *et al*. 2015). This persistent lump was associated with an insufficiently robust instrument that left a fragment of the tag embedded in the whale for nearly 10 years. Similar swellings have been observed in North Atlantic right whales and gray whales tagged with older versions of Type-C tags (Weller 2008; Norman *et al*. 2018). These observations reinforce the need to avoid potential tag breakage in the future.

The lack of significant cyamid infestation of these tag sites, compared to infestation of wounds associated with some entanglement injuries (Rolland *et al*. 2016), suggest there may be minimal necrosis of epithelium at these tag sites, as cyamids feed on shed epithelial cells (Schell *et al*. 2000). A southern right whale seen with a broken tag 11 years after deployment had no significant tissue response around the tag, the tag being observed at the base of a shallow depression in the skin (Best *et al*. 2015). Fragments of a consolidated tag in the blubber of a gray whale were surrounded by epithelial cells with no external observable swelling (Goley *et al*. 2023). Harpoon heads have been observed in the blubber of bowhead whales with minimal tissue response years or decades after embedment, indicating that foreign bodies can embed in epidermal tissues of baleen whales without obvious health impacts (George *et al*. 1999).

Another important predictor of tissue responses to tags, other than tag type, was the placement of the tag on the body. Dorsal application anterior to the dorsal fin was associated with the least swelling. More deeply implanted tags, reflecting a shallower angle of insertion, were associated with less skin loss. Lower photographic scores following tagging were associated with less detectable marks on the skin of the whale years after tagging. Based on these results, every effort should be made to target the dorsum in the vicinity of the dorsal fin, both for the optimization of the research and the welfare of the tagged individual. Experienced taggers and vessel operators are therefore critical for optimal deployments, as are equipment that increases deployment precision, such as tagging platforms and properly-calibrated rifle aims.

Despite the variation in intensity and duration of tissue responses after tagging, most whales had minimal long lasting changes in the skin surface years after tagging. At the end of this study, in some cases, the depression in the skin where the tag had been applied was not detectable, in some others the skin depressions were detectable to the experienced observer with knowledge of the earlier tag site but not to an uninformed observer, and in other whales skin depressions that were detectable were slightly larger than 2–4 times the tag diameter.

In the one whale (Tag 79) with two depressions either side of the dorsal fin, the position of the two depressions, and the larger size of the one contralateral to the tag application site, suggest the tag might have migrated through the whale to exit on the opposite side from tag entry, although no such migration was observed. This is possible, although unlikely, as the tag is fitted with a stopper preventing inward migration. In gunshot animals, bullet exit holes are larger than entry holes, with everted skin margins (Moore *et al*. 2013; Harnish *et al*. 2019). Alternatively, the contralateral lesion could have resulted from fluid discharge tracking along the fascia, exiting the body contralaterally.

The impacts on welfare, as a consequence of pain, are not possible to assess from these photographic data, and remain subjective. Future research on tag effects could focus on methods to evaluate potential for pain assessment, such as changes in behaviour, and stress hormone levels in blow, blubber, baleen or faeces (Hunt *et al*. 2014a; Hunt *et al*. 2014b; Hunt *et al*. 2017; Hunt *et al*. 2019). In addition, continued collaboration amongst the whale research and health monitoring community is vital to ensure all known tagged whales are examined if a carcass should be found, even if years after tagging. Tissues like baleen hold years of whale history and could shed light on stress hormones, reproduction and even feeding ecology changes in tagged whales (Gabriele *et al*. 2021). Concurrently, efforts should continue to improve tag designs to prevent tag breakage and fragment retention and decrease the length and/or diameter of the attachment components to minimise tissue reaction without reducing duration/efficacy. Advanced tag features such as antimicrobial coatings may potentially enhance biocompatibility and thereby increase retention while minimising adverse impacts on the host (Smies *et al*. 2022). The on-going efforts of regional stranding networks are also critical to understanding tagging impacts, which further depends on good communication between those networks and both tagging programs and population studies. Although one tagged whale in this study was found dead following a life-threatening entanglement, no necropsy was possible, the tag wound was not observed, and the individual was also not successfully linked to its life history records until three years later. As a result, there are no relevant observations or samples to further clarify the impacts of the tag. It is further important to note that the tag site of an individual could not necessarily be assessed in all sightings, including, in some cases, its most recent sighting year. Therefore, tag site re-sight data do not represent whale survival, which is presented separately in Robbins *et al*. (in prep).

In conclusion, tag deployments on 79 whales with up to a decade of post-tagging observations revealed that despite a range of local tissue responses mostly influenced by tag design leading to breakage, and position of insertion, the long-term effects detectable at the tag site were minimal, with skin healing and scar contraction. This study led to improved tag designs with robust components, reducing local trauma to tissue resulting from retained tag fragments (Zerbini *et al*., in prep.). It also enabled recognition of the sites on the body least prone to tissue reactions to tag insertion. The decade of observations highlights the value of long-term studies on individually identifiable whales to not only improve understanding of whale ecology, but also to evaluate research methods and their impacts on individual animals. The photographic evaluation scores have contributed to international guidelines for best practices in cetacean tagging (Andrews *et al*. 2019). These results inform researchers as they work to meet ethical and animal welfare standards for research on individual animals while providing valuable information to inform population level conservation.

## ACKNOWLEDGEMENTS

Funding for this project was provided by the National Oceanographic and Atmospheric Administration and ExxonMobil Exploration Company via the National Fish and Wildlife Foundation (NFWF) and the National Oceanographic Partnership Program (NOPP). Support also came from the Marine Mammal Commission and the U.S. Office of Naval Research (ONR, award numbers N-0014-13-1-0653, N-0014-18-1-2749, and N-0014-20-1-2652). The following CCS staff provided assistance in the field: L. Crowe, L. Ganley, T. Kirchner, B. Lynch, E. Sacrey and L. Sette. We also thank Wildlife Computers for working with us to improve the tags over the course of the project. The Gulf of Maine whale watching community contributed data and images of tagged whales and we particularly thank the staff and naturalists of Blue Ocean Society, Boston Harbor Cruises, the Dolphin Fleet, Hyannis Whale Watcher Cruises, New England Coastal Wildlife Alliance, Whale and Dolphin Conservation and the Whale Center of New England. Sightings outside of the Gulf of Maine were shared by the North Atlantic Humpback Whale Catalogue (Allied Whale, College of the Atlantic), R. Etcheberry, J. Frediani and M. MacKay. We thank P. Palsbøll and the students and staff at the University of Groningen for molecular genetic sexes. K. Colegrove conducted a histological analysis of a biopsy sample. The Greater Atlantic Marine Mammal Stranding Network, the Atlantic Marine Conservation Society, the International Fund for Animal Welfare and Virginia Aquarium provided regional stranding information. Thanks to E. King and S. Whiteside from the Australian Antarctic Division for their contributions to the development of Type-C tags for baleen whales. Research was performed under NMFS permits 14245, 20465, 21485, 16325 and 633-1778, the Canadian Department of Fisheries and Oceans as well as IACUC approval from NOAA’s Marine Mammal Laboratory.

## SUPPLEMENTARY MATERIALS

### Supplementary Material 1 - Photogrammetry

A photogrammetric method was used to measure features of interest in follow-up photographs. Initially, a range finder was used in the field to measure the distance from the camera to the whale (e.g., Jaquet 2006). However, the Jaquet (2006) approach limited measurements to photographs with paired range finder data and also required that each camera/lens combination be calibrated across the relevant subject distances and focal lengths. This study used images from 21 different camera models, including contributions from opportunistic observers. An alternative measurement method was therefore used that took advantage of the fact that an object of known size (the tag) had been placed on the whale.

Photogrammetric measurements were conducted for all whales with suitable photographic coverage. Known sized objects were measured using the program ICY (de Chaumont *et al*. 2012) to establish a scale (mm/pixel) for objects at approximately the same distance in each image. The tag was initially used as the scale, but we also measured persistent natural features on the whale to act as reference marks when the tag was no longer present.

Two reference marks were chosen for each tagged whale. These marks were selected based on their persistence across years, their proximity to the tag site and their size. The two marks selected were typically substantially different in size and as orthogonal to each other as possible. Measurements were made on the subset of photos that were taken approximately perpendicular to the feature of interest, which was either along the long axis (i.e., anterior/posterior axis) of the whale, or directly behind it. The distance from the camera to the whale was typically much greater than the size of the objects being measured and therefore slight deviations from the preferred angle (i.e., within 15°) were expected to produce minimal error. A direct comparison of this approach to the Jaquet (2006) method suggested comparable accuracy, but greater ease of use and wider applicability to the data in this study.The maximum depth of tag penetration was calculated based on a measurement of the minimum length of tag exposed on the day of deployment, the insertion angle and the known length of the tag (Figure 8). It was calculated as follows:

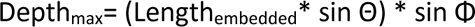

where: Length_embedded_ = Length_total tag_ – Length_exposed_, Θ = angle in the dorsal-ventral plane and Φ = angle in the anterior-posterior plane.

Angles were measured using the Angle Helper tool in program ICY. Photos taken perpendicular the dorsal-ventral and the anterior-posterior planes were used to calculate the angle incident to each. These were selected such that the view of the surface of the skin surface at the point of tag insertion was approximately normal to the camera lens. An anterior-posterior angle was also estimated assuming that the measured length of the exposed tag was one length of a right-angle triangle and the slope from the base of the tag to the top, as viewed from above the tag, was the opposite length. When possible, this second method was used as a check of anterior-posterior angle calculations.

**Fig. S1:**
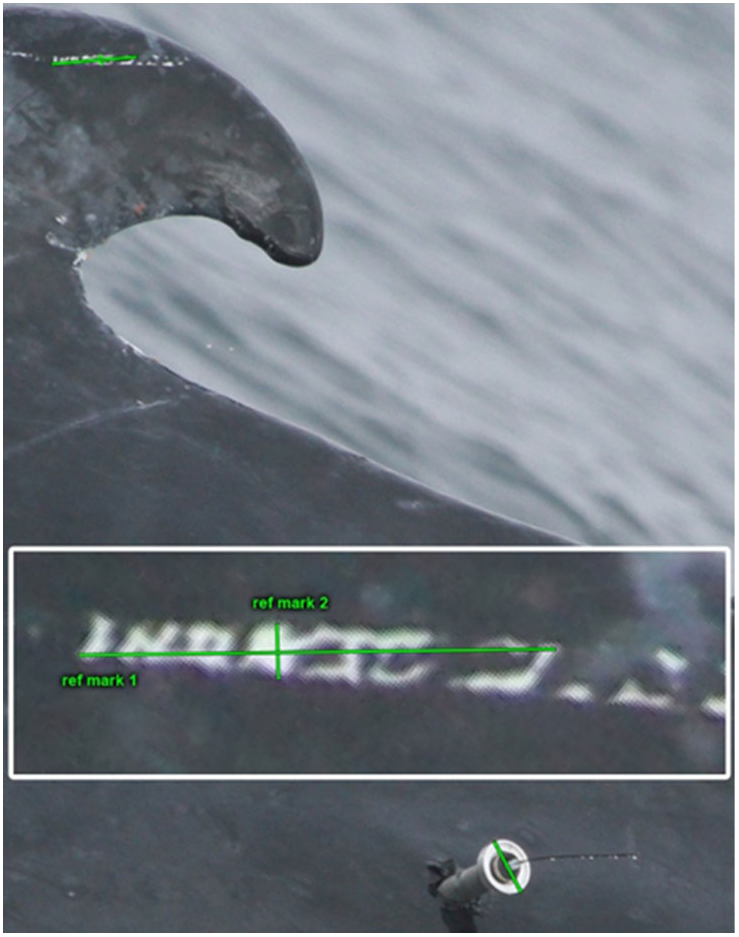
Example of reference marks established for photogrammetry. Natural marks on the dorsal fin were measured while a tag of known size was present to establish a scale. The inset shows greater detail of the tag and natural markings that were measured. Image credit: CCS.

**Fig. S2:**
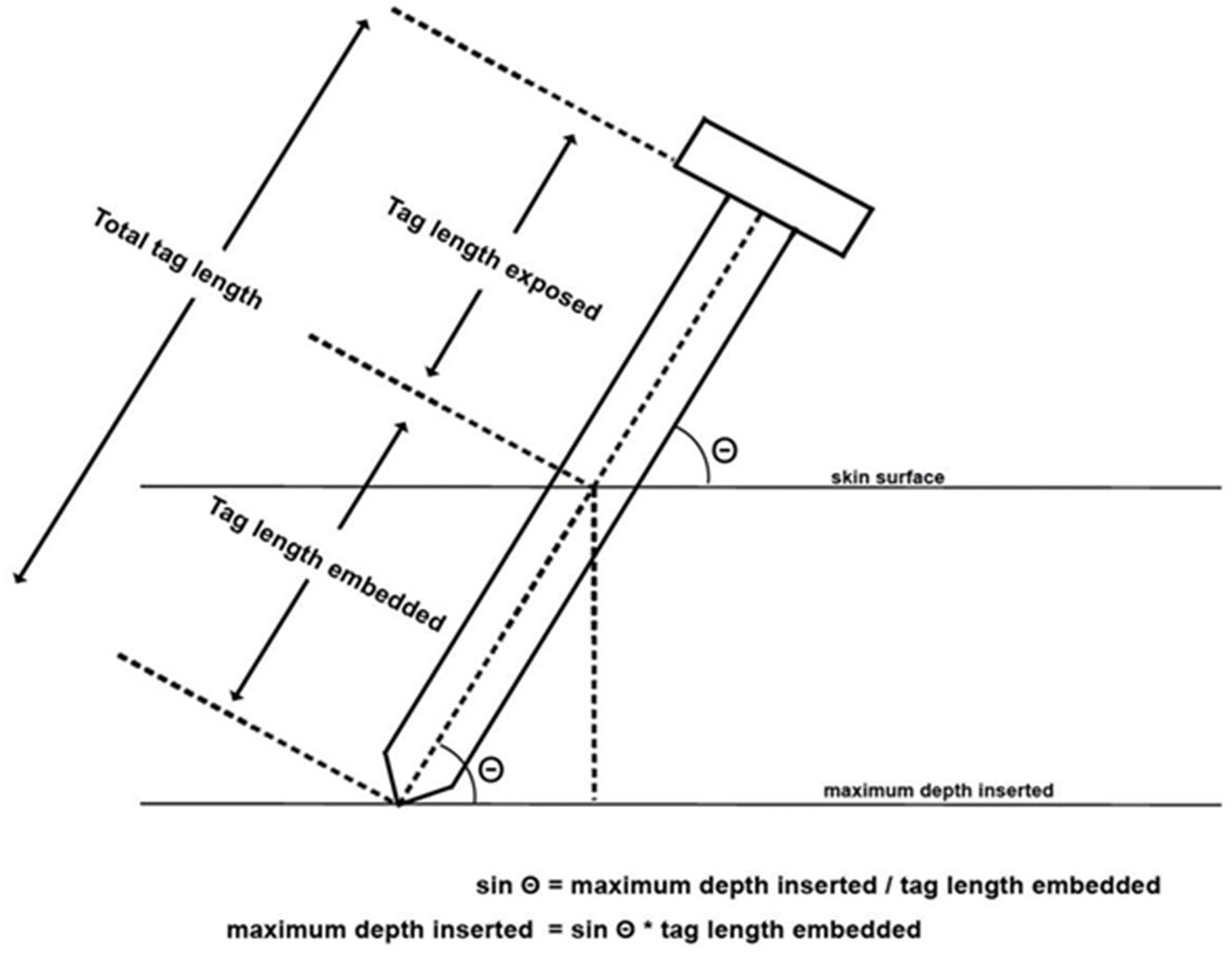
Schematic of measures used to estimate maximum penetration depth.

### Supplementary Material 2 - Results

**Table S1.** Summary of sample sizes used for statistical analyses according to whale sex, year of tag deployment, tag design and tag placement on the body.

| Variable | Individuals | Sightings |
| --- | --- | --- |
| Sex |  |  |
| Female | 35 | 1204 |
| Male | 36 | 1764 |
| Unknown | 1 | 3 |
| Year of tag deployment |  |  |
| 2011 | 18 | 1534 |
| 2012 | 11 | 414 |
| 2013 | 7 | 260 |
| 2015 | 21 | 540 |
| 2018 | 15 | 223 |
| Tag design |  |  |
| Integrated | 43 | 1023 |
| Non-integrated | 29 | 1948 |
| Tag position |  |  |
| Dorsal – Cranial | 20 | 804 |
| Dorsal – Central | 30 | 1463 |
| Ventral – Cranial | 5 | 127 |
| Ventral – Central | 17 | 577 |
| **Total** | **72** | **2971** |

**Table S2.**
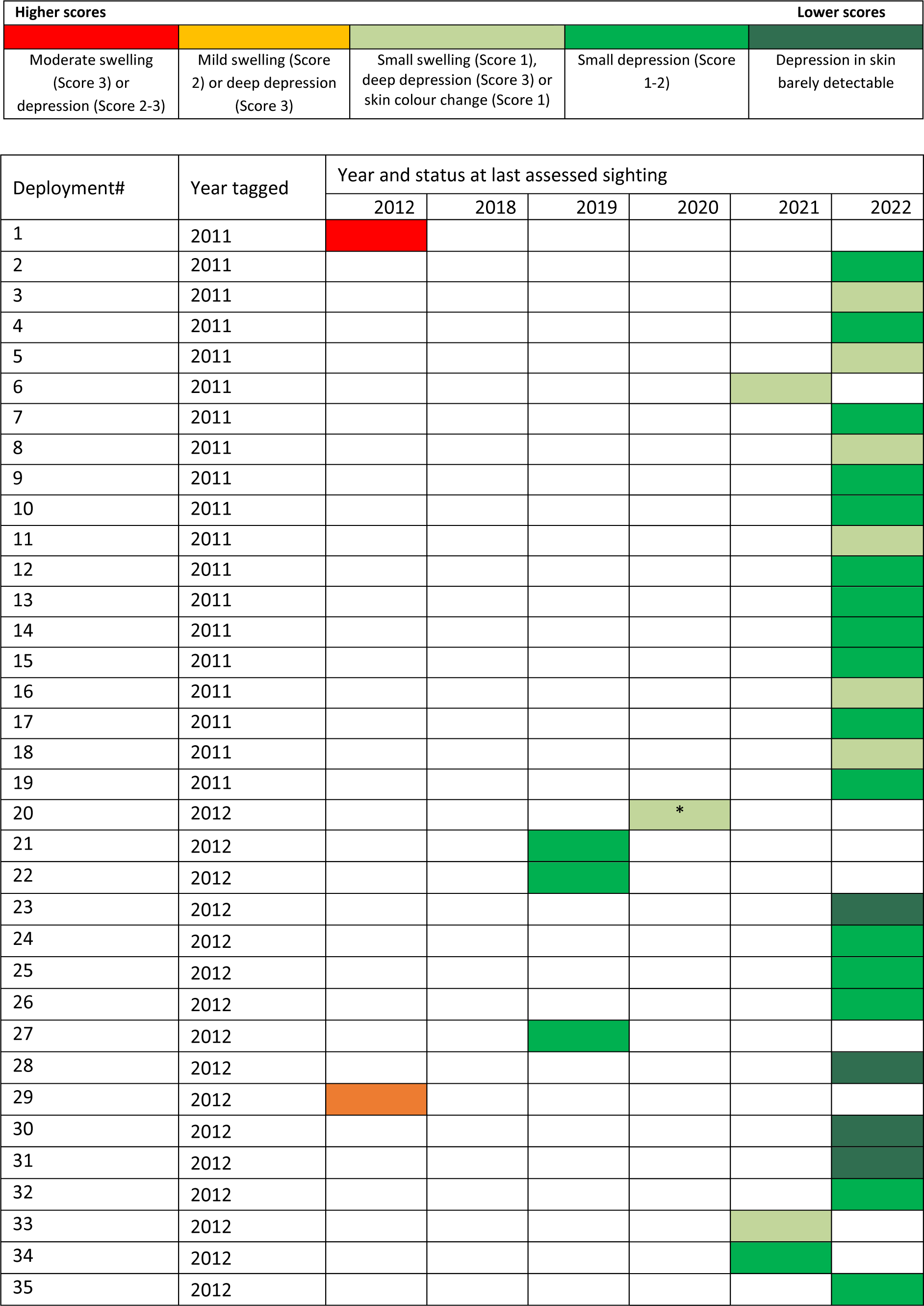
Status of the tag site at last photographed sighting for whales tagged with non-integrated tags. Colour codes are as shown below. Note: *= Individual last seen in poor health.

**Table S3.**
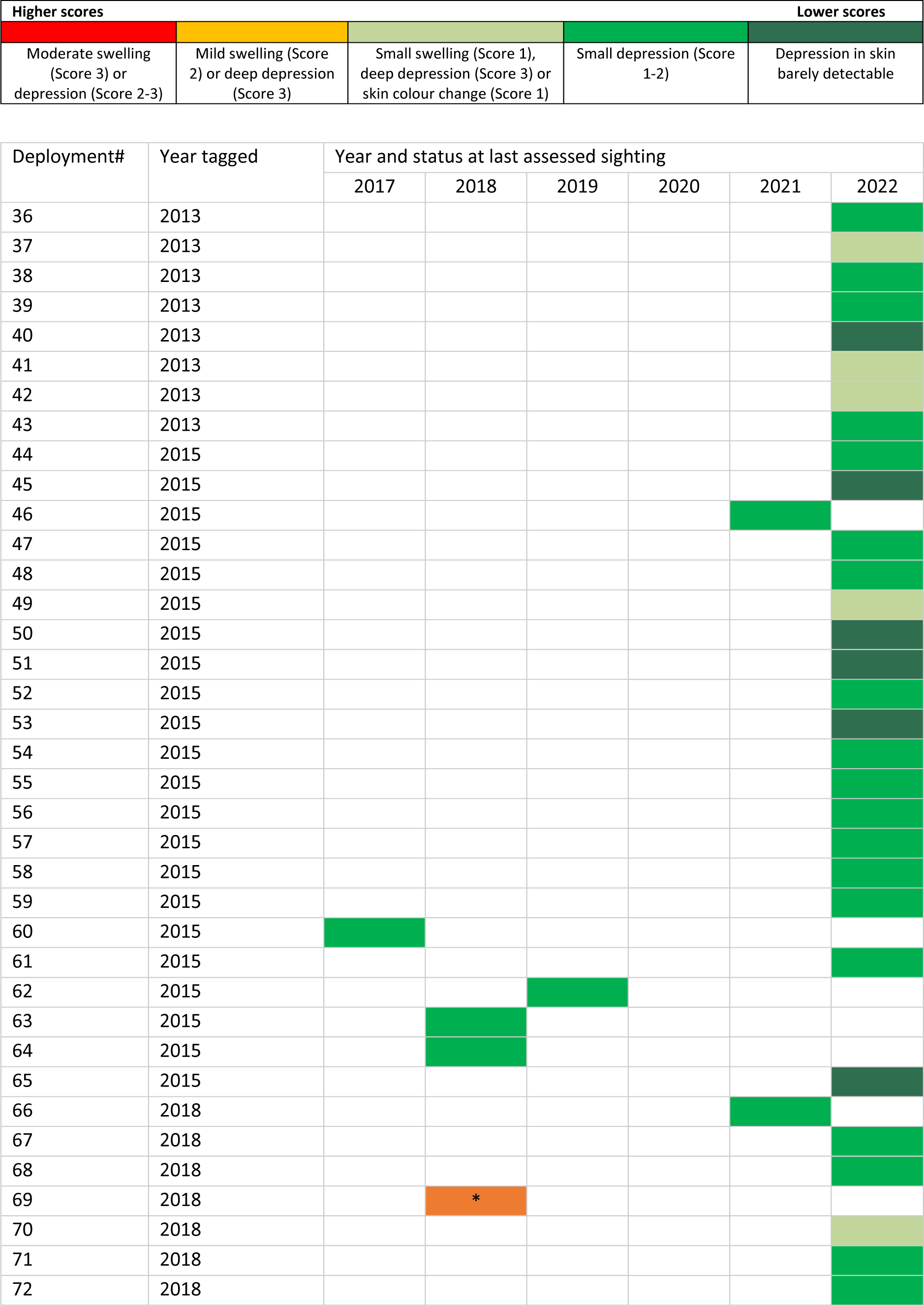
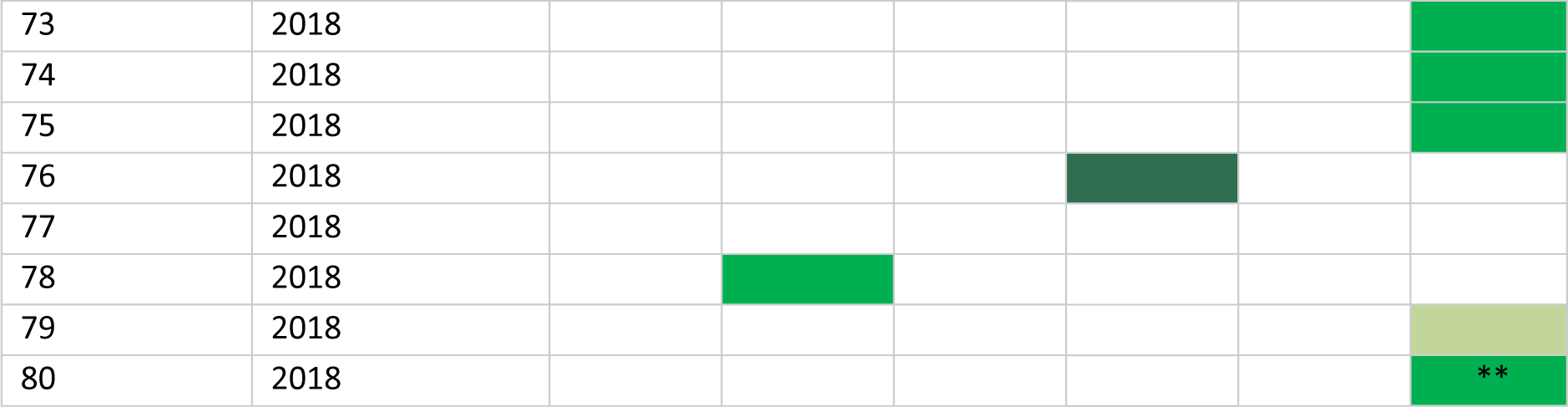
Status of the tag site at last photographed sighting for whales tagged with integrated tags. Colour codes are as shown below. Notes: *=This individual died in 2019, but was involved in a life-threatening entanglement event that dislodged the tag 10 days after tagging. **=There was no adequate tag site image in 2022, so an image from 2023 was analysed.

## REFERENCES

Andrews R., Baird R. W., Calambokidis J., Goertz C. E. C., Gulland F. M. D., Heide-Jørgensen M. P., Hooker S. K., Johnson M., Mate B., Mitani Y., Nowacek D. P., Owen K., Quakenbush L. T., Raverty S., Robbins J., Schorr G. S., Shpak O. V., Townsend J., F.I., Uhart M., Wells R. S., Zerbini A. N. 2019. Best Practice Guidelines for Cetacean Tagging. Journal of Cetacean Research and Management 20:27–66.

Bailey I. S., Karran S. E., Toyn K., Brough P., Ranaboldo C., Karran S. J. 1992. Community surveillance of complications after hernia surgery. British Medical Journal 304(6825):469–471.

Bates D., Mächler M., Bolker B., Walker S. 2015. Fitting Linear Mixed-Effects Models Using lme4. Journal of Statistical Software 67(1):1–48.

Baumgartner M. F., Mate B. R. 2005. Summer and fall habitat of North Atlantic right whales (*Eubalaena glacialis*) inferred from satellite telemetry. Canadian Journal of Fisheries and Aquatic Sciences 62:527–543.

Bérubé M., Palsbøll P. J. 1996a. Identification of sex in cetaceans by multiplexing with three ZFX and ZFY specific primers. Molecular Ecology 5:283–287.

Bérubé M., Palsbøll P. J. 1996b. Identification of sex in cetaceans by multiplexing with three ZFX and ZFY specific primers: erratum. Molecular Ecology 5:602.

Best P., Mate B. 2007. Sighting history and observations of southern right whales following satellite tagging off South Africa. Journal of Cetacean Research and Management 9:111–114.

Best P. B., Mate B., Lagerquist B. 2015. Tag retention, wound healing, and subsequent reproductive history of southern right whales following satellite-tagging. Marine Mammal Science 31(2):520–539.

Bradford A. L., Weller D. W., Punt A. E., Ivashchenko Y. V., Burin A. M., VanBlaircom G. R., Brownell Jr R. L. 2012. Leaner leviathans: body condition variation in a critically endangered whale population. Journal of Mammalogy 93:251–266.

Bruce-Allen L. J., Geraci J. R. 1985. Wound-healing in the bottlenose dolphin (*Tursiops truncatus*). Canadian Journal of Fisheries and Aquatic Sciences 42(2):216–228.

Burek-Huntington K. A., Shelden K. E. W., Andrews R. D., Goertz C. E. C., McGuire T. L., Dennison S. 2023. Postmortem pathology investigation of the wounds from invasive tagging in belugas (*Delphinapterus leucas*) from Cook Inlet and Bristol Bay, Alaska. Marine Mammal Science 39(2):492–514.

Charlton C., Christiansen F., Ward R., Mackay A. I., Andrews-Goff V., Zerbini A. N., Childerhouse S., Guggenheimer S., O’Shannessy B., Brownell Jr R. L. 2023. Evaluating short to medium term effects of implantable satellite tags on southern right whales *Eubalaena australis*. Diseases of Aquatic Organisms 155:125–140.

Christensen R. 2022. *ordinal*: Regression Models for Ordinal Data. R package version 2022.11-16, https://CRAN.R-project.org/package=ordinal.

Christiansen F., Sironi M., Moore M. J., Di Martino M., Ricciardi M., Warick H. A., Irschick D. J., Gutierrez R., Uhart M. M. 2019. Estimating body mass of free-living whales using aerial photogrammetry and 3D volumetrics. Methods in Ecology and Evolution 10(12):2034–2044.

Clapham P. J. 1992. Age at attainment of sexual maturity in humpback whales, *Megaptera novaeangliae*. Canadian Journal of Zoology 70:1470–1472.

Clapham P. J., Baraff L. S., Carlson C. A., Christian M. A., Mattila D. K., Mayo C. A., Murphy M. A., Pittman S. 1993. Seasonal occurrence and annual return of humpback whales, *Megaptera novaeangliae*, in the southern Gulf of Maine. Canadian Journal of Zoology 71:440–443.

de Chaumont F., Dallongeville S., Chenouard N., Herve N., Pop S., Provoost T., Meas-Yedid V., Pankajakshan P., Lecomte T., Le Montagner Y., Lagache T., Dufour A., Olivo-Marin J.-C. 2012. Icy: an open bioimage informatics platform for extended reproducible research. Nat Meth 9(7):690–696.

Fox J., Weisberg S., Price B., Friendly M., Hong J., Anderson R., Firth D., Taylor S., Core Team R. 2022. *effects*: Effect Displays for Linear, Generalized Linear, and Other Models. R package version 4.2-2. https://cran.r-project.org/web/packages/effects/index.html.

Gabriele C. M., Taylor L. F., Huntington K. B., Buck C. L., Hunt K. E., Lefebvre K. A., Lockyer C., Lowe C., Moran R., Murphy A., Rogers M. C., Trumble S. J., Raverty S. 2021. Humpback whale #441 (Festus): Life, death, necropsy, and research findings. Natural Resource Report NPS/GLBA/NRR—2021/2250. National Park Service, Fort Collins, Colorado. 10.36967/nrr-2285345.

Gales N., Double M. C., Robinson S., Jenner C., Jenner M., King E., Gedamke J., Paton D., Raymond B. 2009a. Satellite tracking of southbound East Australian humpback whales (*Megaptera novaeangliae*); challenging the feast or famine model for migrating humpback whales. Paper SC/61/SH17 presented to the IWC Scientific Committee, 31 May-12 June 2009, Funchal, Madeira, Portugal (unpublished). 12pp. [Paper available from the Office of this Journal].

Gales N. J., Bowen W. D., Johnston D. W., Kovacs K. M., Littnan C. L., Perrin W. F., Reynolds III J. E., Thompson P. M. 2009b. Guidelines for the treatment of marine mammals in field research. Marine Mammal Science 25(3):725–736.

Gendron D., Serrano I. M., de la Cruz A. U., Calambokidis J., Mate B. 2015. Long-term individual sighting history database: an effective tool to monitor satellite tag effects on cetaceans. Endangered Species Research 26(3):235–241.

George J. C., Bada J., Zeh J., Scott L., Brown S. E., O’Hara T., Suydam R. 1999. Age and growth estimates of bowhead whales (*Balaena mysticetus*) via aspartic acid racemization. Canadian Journal of Zoology 77:571–580.

Glockner D. A. 1983. Determining the sex of humpback whales in their natural environment. Pages 447-464 in Payne R, editor. Communication and Behavior of Whales. AAAS Selected Symposium 76. Westview Press, Colorado.

Goley P. D., Calambokidis J., Duignan P., Gulland F. M. D., Halaska B., Lui A., Martinez M., Mate B. 2023. Observations of tissue healing around an implanted “C” tag in a Pacific Coast Feeding Group gray whale (*Eschrichtius robustus*). Marine Mammal Science 40(1):254–261.

Gorad K., Rahate V., Shinde G., Taralekar G., Prabhu V., Singh L. 2021. Use of Southampton Scoring for Wound Healing in Post-surgical Patients: Our Experience in Semi-urban Setup. Archives of Clinical and Biomedical Research 5:36–41.

Gulland F. M. D., Dierauf L., Whitman K. e., editors. 2018. CRC Handbook of Marine Mammal Medicine, Third edition, CRC Press, Boca Raton, FL, 1063 pp.

Harnish A. E., Ault J., Babbitt C., Gulland F. M. D., Johnson P. C., Shaughnessy N. L., Wood K. A., Baird R. W. 2019. Survival of a Common Bottlenose Dolphin (Tursiops truncatus) Calf with a Presumptive Gunshot Wound to the Head. Aquatic Mammals 45(5):543–548.

Heide-Jørgensen M. P., Kleivane L., Oien N., Laidre K. L., Jensen M. V. 2001. A new technique for deploying satellite transmitters on baleen whales: Tracking a blue whale (*Balaenoptera musculus*) in the North Atlantic. Marine Mammal Science 17(4):949–954.

Hill A., Karniski C., Robbins J., Pitchford T., Todd S., Asmutis-Silvia R. 2017. Vessel collision injuries on live humpback whales, *Megaptera novaeangliae*, in the southern Gulf of Maine. Marine Mammal Science 33:558–573.

Hörbst S. 2019. Visual health assessment of parous female southern right whales (*Eubalaena australis*) off the southern Cape coast, South Africa. Master’s Thesis. Faculty of Science, Department of Biological Sciences. 49pp. http://hdl.handle.net/11427/31584. pp.

Hunt K. E., Lysiak N. S., Robbins J., Moore M. J., Seton R. E., Torres L., Buck C. L. 2017. Multiple steroid and thyroid hormones detected in baleen from eight whale species. Conservation Physiology 5(1):cox061.

Hunt K. E., Robbins J., Buck C. L., Bérubé M., Rolland R. M. 2019. Evaluation of fecal hormones for noninvasive research on reproduction and stress in humpback whales (*Megaptera novaeangliae*). General and Comparative Endocrinology 280:24–34.

Hunt K. E., Rolland R. M., Kraus S. D. 2014a. Detection of steroid and thyroid hormones via immunoassay of North Atlantic right whale ( Eubalaena glacialis) respiratory vapor. Marine Mammal Science 30(2):796–809.

Hunt K. E., Stimmelmayr R., George C., Hanns C., Suydam R., Brower H., Rolland R. M. 2014b. Baleen hormones: a novel tool for retrospective assessment of stress and reproduction in bowhead whales (*Balaena mysticetus*). Conservation Physiology 2(1):cou030.

Jaquet N. 2006. A simple photogrammetric technique to measure sperm whales at sea. Marine Mammal Science 22:862–879.

Katona S. K., Whitehead H. P. 1981. Identifying humpback whales using their natural markings. Polar Record 20:439–444.

Knowlton A. R., Hamilton P. K., Marx M. M., Pettis H. M., Kraus S. D. 2012. Monitoring North Atlantic right whale *Eubalaena glacialis* entanglement rates: a 30 yr retrospective. Marine Ecology Progress Series 466:293–302.

Kraus S., Quinn C., Slay C. 2000. Report on the workshop on the effects of tagging on North Atlantic right whales. New England Aquarium, Central Wharf, Boston, Massachusetts. October 23, 1999.

Lockyer C. 1981. Growth and energy budgets of large baleen whales from the Southern Hemisphere. FAO Fisheries Series No. 5 3:397–487.

Mate B., Mesecar R., Lagerquist B. 2007. The evolution of satellite-monitored radio tags for large whales: One laboratory’s experience. Deep-Sea Research Part II-Topical Studies in Oceanography 54(3-4):224-247.

Mate B. R., Harvey J. T., Hobbs L., Maiefski R. 1983. A new attachment device for radio-tagging large whales. Journal of Wildlife Management 47:868–872.

Mehta A. V., Allen J. M., Constantine R., Garrigue C., Jann B., Jenner C., Marx M. K., Matkin C. O., Mattila D. K., Minton G., Mizroch S. A., Olavarria C., Robbins J., Russell K. G., Seton R. E., Steiger G. H., Vikingsson G. A., Wade P. R., Witteveen B. H., Clapham P. J. 2007. Baleen whales are not important as prey for killer whales (*Orcinus orca*) in high latitudes. Marine Ecology Progress Series 348:297–307.

Minton G., Van Bressem M. F., Willson A., Collins T., Harthis A. L., Sarrouf Willson M., Baldwin R., Leslie M., van Waerbeek K. 2022. Visual health assessment and evaluation of anthropogenic threats to Arabian Sea humpback whales in Oman. Journal of Cetacean Research and Management 23:59–79.

Mizroch S. A., Tillman M. F., Jurasz S., Straley J. M., von Ziegesar O., Herman L. M., Pack A. A., Baker S., Darling J., Glockner-Ferrari D., Ferrari M., Salden D. R., Clapham P. J. 2011. Long-term survival of humpback whales radio-tagged in Alaska from 1976 through 1978. Marine Mammal Science 27(1):217–229.

Moore M., Andrews R., Austin T., Bailey J., Costidis A., George C., Jackson K., Pitchford T., Landry S., Ligon A., McLellan W., Morin D., Smith J., Rotstein D., Rowles T., Slay C., Walsh M. 2013. Rope trauma, sedation, disentanglement, and monitoring-tag associated lesions in a terminally entangled North Atlantic right whale (*Eubalaena glacialis*). Marine Mammal Science 29(2):E98–E113.

Moore M., Zerbini A. 2017. Dolphin blubber/axial muscle shear: implications for rigid transdermal intramuscular tracking tag trauma in whales. Journal of Experimental Biology 220(20):3717–3723.

Murray J. C., Pollack S. V., Pinnell S. R. 1981. Keloids - a review. Journal of the American Academy of Dermatology 4(4):461–470.

Naessig P. J., Lanyon J. M. 2004. Levels and probable origin of predatory scarring on humpback whales (*Megaptera novaeangliae*) in east Australian waters. Wildlife Research 31:163–170.

Nickoloff B. J., Naidu Y. 1994. Perturbation of epidermal barrier function correlates with initiation of cytokine cascade in human skin. Journal of the American Academy of Dermatolology 30(4):535–546.

Norman S. A., Flynn K. R., Zerbini A. N., Gulland F. M. D., Moore M. J., Raverty S., Rotstein D. S., Mate B. R., Hayslip C., Gendron D., Sears R., Douglas A. B., Calambokidis J. 2018. Assessment of wound healing of tagged gray (*Eschrichtius robustus*) and blue (*Balaenoptera musculus*) whales in the eastern North Pacific using long-term series of photographs. Marine Mammal Science 34(1):27–53.

ONR. 2009. Final Workshop Proceedings of the Cetacean Tag Design Workshop. Office of Naval Research, Arlington, Virginia. 18pp.

Palsbøll P. J., Larsen F., Sigurd Hansen E. 1991. Samping of skin biopsies from free-ranging large cetaceans in West Greenland: Development of new biopsy tips and bolt designs. Reports of the International Whaling Commission (special issue) 13:71–79.

Palsbøll P. J., Vader A., Bakke I., El-Gewely M. R. 1992. Determination of gender in cetaceans by the polymerase chain reaction. Canadian Journal of Zoology 70:2166–2170.

Papastavrou V., Ryan C. 2023. Ethical standards for research on marine mammals. Research Ethics 19(4):390–408.

Pettis H. M., Rolland R. M., Hamilton P. K., Brault S., Knowlton A. R., Kraus S. D. 2004. Visual health assessment of North Atlantic right whales (*Eubalaena glacialis*) using photographs. Canadian Journal of Zoology 82(1):8–19.

R Core Team. 2012.R: A language and environment for statistical computing.

Ripley B., Venables B., Bates D. M., Hornik K., Gebhardt A., Firth D. 2022. *Mass*: Support Functions and Datasets for Venables and Ripley’s MASS. R package version 7.3-60. https://cran.r-project.org/web/packages/MASS/index.html.

Robbins J. 2007. Structure and dynamics of the Gulf of Maine humpback whale population. Ph.D. thesis. St Andrews, Scotland. 179 pp.

Robbins J., Mattila D. K. 2004. Estimating humpback whale (Megaptera novaeangliae) entanglement rates on the basis of scar evidence. Report to the National Marine Fisheries Service. Order number 43ENNF030121. 22pp.

Robbins J., Zerbini A. N., Gales N., Gulland F. M. D., Double M., Clapham P. J., Andrews-Goff V., Kennedy A. S., Landry S., Mattila D. K., Tackaberry J. 2013. Satellite tag effectiveness and impacts on large whales: preliminary results of a case study with Gulf of Maine humpback whales. Paper SC/65a/SH05 presented to the IWC Scientific Commitee, 3-15 June 2013, Jeju, South Korea (unpublished). 10pp [Paper available from the Office of this Journal]

Rolland R. M., Schick R. S., Pettis H. M., Knowlton A. R., Hamilton P. K., Clark J. S., Kraus S. D. 2016. Health of North Atlantic right whales Eubalaena glacialis over three decades: from individual health to demographic and population health trends. Marine Ecology Progress Series 542:265–282.

Schell D. M., Rowntree V. J., Pfeiffer C. J. 2000. Stable-isotope and electron-microscopic evidence that cyamids (Crustacea : Amphipoda) feed on whale skin. Canadian Journal of Zoology 78(5):721–727.

Sepúlveda M., Perez-Alvarez M. J., Santos-Carvallo M., Pavez G., Olavarria C., Moraga R., Zerbini A. N. 2018. From whaling to whale watching: Identifying fin whale critical foraging habitats off the Chilean coast. Aquatic Conservation-Marine and Freshwater Ecosystems 28(4):821–829.

Slijper E. J. 1962. Whales. Cornell University Press, Ithaca, New York. p 299.

Smies A., Wales J., Hennenfent M., Lyons L., Dunn C., Robbins J., Lee B. P., Zerbini A., Rajachar R. M. 2022. Non-antibiotic antimicrobial polydopamine surface coating to prevent stable biofilm formation on satellite telemetry tags used in cetacean conservation applications. Frontiers in Marine Science 9.

Steiger G. H., Calambokidis J., Straley J. M., Herman L. M., Cerchio S., Salden D. R., Urban-R J., Jacobsen J. K., von Ziegesar O., Balcomb K. C., Gabriele C. M., Dahlheim M. E., Uchida S., Ford J. K. B., Ladron de Guevara-P P., Yamaguchi M., Barlow J. 2008. Geographic variation in killer whale attacks on humpback whales in the North Pacific: implications for predation pressure. Endangered Species Research 4:247–256.

Watkins W. A., Moore K. E., Wartzok D., Johnson J. H. 1981. Radio tracking of finback (*Balaenoptera physalus*) and humpback (*Megaptera novaeangliae*) whales in Prince William Sound, Alaska. Deep Sea Research Part A. Oceanographic Research Papers 28(6):577–588.

Weller D. W. 2008. Report of the Large Whale Tagging Workshop convened by the U.S. Marine Mammal Commission and the U.S. National Marine Fisheries Service, San Diego, California. 32pp.

Wickham H. 2016. ggplot2: Elegant Graphics for Data Analysis. Springer-Verlag New York. ISBN 978–3-319-24277-4, https://ggplot2.tidyverse.org.

Wickham H. 2022. *plyr*: Tools for Splitting, Applying and Combining Data. R package version 1.8.8. https://cran.r-project.org/web/packages/plyr/index.html.

Wilke C. 2020. *cowplot*: Streamlined Plot Theme and Plot Annotations for ‘ggplot2’. R package version 1.1.1, https://wilkelab.org/cowplot/.

Willson A., Leslie M., Baldwin R., Cerchio S., Childerhouse S., Collins T., Findlay K., Genov T., Godley B. J., Al Harthi S., Macdonald D. W., Minton G., Zerbini A. N., Witt M. J. 2018. Update on satellite telemetry studies and first unoccupied aerial vehicle assisted health assessment studies of Arabian Sea humpback whales off the coast of Oman. Paper IWC/SC67B/CMP13Rev1 presented to the IWC Scientific Commitee, 24 April-6 May 2018, Bled, Slovenia (unpublished). 14pp. [Paper available from the Office of this Journal].

Womersley F., Hancock J., Perry C. T., Rowat D. 2021. Wound-healing capabilities of whale sharks (*Rhincodon typus*) and implications for conservation management. Conservation Physiology 9:16.

Zasloff M. 2011. Observations on the Remarkable (and Mysterious) Wound-Healing Process of the Bottlenose Dolphin. Journal of Investigative Dermatology 131(12):2503–2505.

Zerbini A. N., Ajo, A.F., Andriolo, A., Clapham, P.J., Crespo, E., Gonzalez, R., Harris, G., Mendez, M., Rosenbaum, H., Sironi, M., Sucunza, F. and Uhart, M. 2018. Satellite tracking of Southern right whales (*Eubalaena australis*) from Golfo San Matías, Rio Negro Province, Argentina. Paper SC/67B/CMP/17 presented to the IWC Scientific Committee, 23 April-6 May 2018, Bled, Slovenia (unpublished) 10pp. [Paper available from the Office of this Journal].

Zerbini A. N., Andriolo A., Heide-Jørgensen M. P., Pizzorno J. L., Maia Y. G., VanBlaricom G. R., DeMaster D. P., Simões-Lopes P. C., Moreira S., Bethlem C. 2006. Satellite-monitored movements of humpback whales *Megaptera novaeangliae* in the Southwest Atlantic Ocean. Marine Ecology Progress Series 313:295–304.

Zerbini A. N., Baumgartner M. F., Kennedy A. S., Rone B. K., Wade P. R., Clapham P. J. 2015. Space use patterns of the endangered North Pacific right whale *Eubalaena japonica* in the Bering Sea. Marine Ecology Progress Series 532:269–281.

